# H3K27 trimethylation maintains baseline network excitability in epilepsy

**DOI:** 10.64898/2026.07.24.740616

**Authors:** Jose Ezekiel C. Espina, Olivia R. Hoffman, Jennifer L. Koehler, Barry A. Schoenike, Avtar S. Roopra

## Abstract

The processes by which epileptic insults precipitate the molecular, cellular, and network alterations in the brain that lead to epilepsy are poorly understood. We previously discovered that after *status epilepticus* (SE – an epilepsy inducing severe bout of seizures) the H3K27 methylase Enhancer of Zeste Homolog 2 (EZH2) is robustly induced and drives repression of genes. Both systemic pharmacological inhibition of EZH2 and deletion of EZH2 in neurons exacerbates epilepsy progression, suggesting that acute EZH2 induction may exert a net protective effect against disease progression chronically. However, the mechanisms underlying EZH2-mediated control of the putative protective and pathological pathways in disease progression are still unknown. To interrogate the mechanisms of EZH2 function post-SE, we used bulk CUT&RUN-sequencing against H3K27me3 in tandem with bulk RNA-sequencing in hippocampi of naive and 4d. post-SE mice to profile epigenomic and transcriptomic changes. Differential peak analysis showed that H3K27me3 was enriched both at loci pre-marked by H3K27me3 in the naive hippocampus as well as in loci that were de novo methylated after SE, consistent with the SE-dependent induction of EZH2 protein levels. Multi-omic integration of CUT&RUN and RNA-seq data revealed a module of genes that were coordinately H3K27me3 enriched, transcriptionally repressed, and annotated to ontological terms involved in neuronal signaling and network excitability. This result suggests that EZH2 induction may function in part to control network excitability after injury. To test this, we treated mice acutely post-SE with the EZH2 inhibitor UNC1999 and found that EZH2 inhibition attenuated H3K27me3 induction and dampened repression of target network excitability genes in response to SE. Functionally, UNC1999 treatment significantly increased seizure probability acutely and exacerbated disease severity in the chronic period. Taken together, these results suggest that EZH2 induction after injury may function to maintain network excitability to lower seizure probability acutely to protect against disease progression.

## INTRODUCTION

Epilepsy is one of the most common neurological disorders and is characterized by the occurrence of spontaneous recurrent seizures [1–3]. While current treatments for epilepsy are effective in controlling seizures for roughly 2/3 of patients, the remaining 1/3 of patients go on to develop drug-refractory epilepsy [4]. Moreover, current treatments do nothing to reverse nor prevent the underlying mechanisms that propagate and maintain the disease thereby necessitating continuous medication for a patient’s lifespan [5]. This lack of disease modifying treatments in epilepsy is due in part to our poor understanding of the suite of molecular, cellular, and network alterations that lead to the occurrence of spontaneous seizures.

The process that transforms a healthy brain to one supporting the generation of spontaneous seizures (ie. an epileptic brain) is known as epileptogenesis. In acquired epilepsies, epileptogenesis is precipitated by an initiating brain insult such as trauma, an infection, or an unremitting bout of seizures known as *status epilepticus* (SE) [6]. SE is commonly defined as a prolonged seizure lasting more than 5 minutes or 2 or more seizures with incomplete recovery of consciousness in between [7–10]. In pre-clinical models, SE can be induced either by electrical or chemical stimulation [11]. SE is then followed by a seizure-free period known as the latent period before the emergence of spontaneous seizures in a period known as the chronic period.

Our lab identified the transcriptional repressor Enhancer of Zeste Homolog 2 (EZH2) as a master regulator of gene repression in the early phase of epileptogenesis where its protein levels are robustly induced acutely post-SE but abate within 10-20 days [12,13]. Deletion in neurons or systemic pharmacologic inhibition of EZH2 during this period exacerbates disease severity in the chronic period, suggesting that EZH2 exerts a net protective effect against disease progression, but the mechanisms are unknown [12,13]. EZH2 is one of the core sub-units of the Polycomb Repressive Complex 2 (PRC2), which functions to silence gene expression through methylation of the histone H3 tail at lysine 27 (H3K27) to facilitate chromatin compaction [14–16]. PRC2 function is essential for embryonic development and maintenance of cell identity [15,17,18], however, its role in epilepsy disease progression is poorly understood.

Here we show that H3K27me3 is induced acutely post-SE consistent with increases in EZH2 protein levels. H3K27me3 deposition is associated with repression of a module of genes controlling neuronal signaling and network excitability. Inhibition of EZH2 activity post-SE attenuates H3K27me3 deposition and diminishes RNA repression at these genes. Functionally, EZH2 inhibition increases network excitability acutely post-SE and is associated with increased seizure frequency chronically.

## RESULTS

### *Status epilepticus* induces H3K27me3 enrichment at pre-marked and de novo loci

EZH2 is induced across cell types after SE [12,13]. To test whether this induction results in altered H3K27 methylation, we performed CUT&RUN-sequencing against H3K27me3 in nuclei isolated from hippocampi of mice 4 days post-SE (termed post-SE from hereon) and in naive controls and quantified changes in H3K27me3 distribution induced by SE (Fig. 1A). Differential enrichment analysis identified 3,442 H3K27me3 peaks that were altered in post-SE hippocampi vs. naive control at an FDR cut-off of 0.05 (Fig. 1B, S1 Data). Though a number of H3K27me3 peaks were depleted post-SE (n=783) (Fig. 1B), the majority (2659 peaks) showed increased H3K27me3 signal post-SE. Heatmaps and metaplot representations of H3K27me3 signal in naive and post-SE hippocampi (Fig. 1C) highlight the broad signal that is characteristic of H3K27me3 peaks [19].

**Fig. 1.**
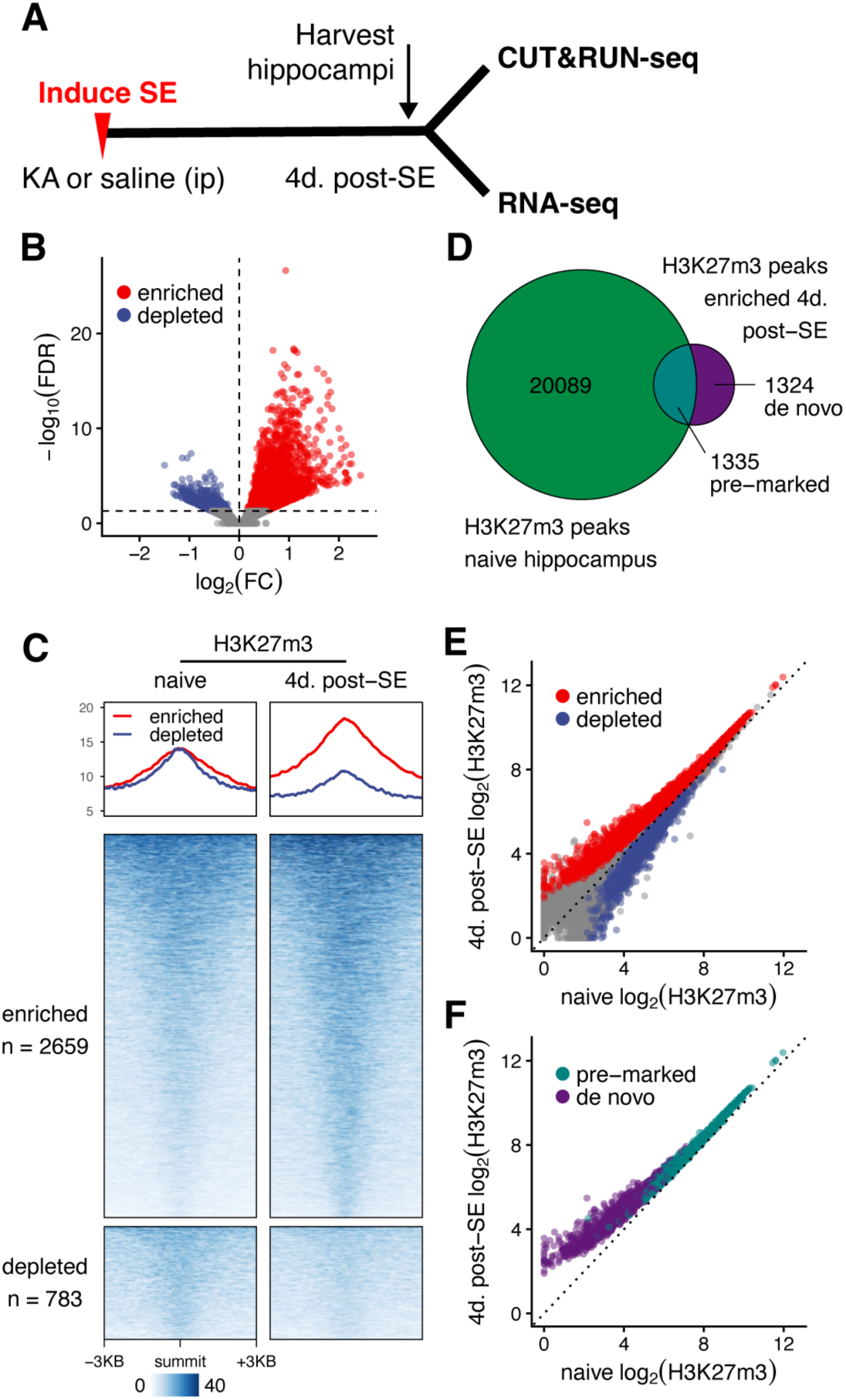
H3K27me3 is enriched 4d. post-SE in pre-marked and de novo loci. **(A)** Experimental workflow. Mice were induced to SE or treated with saline as control. Hippocampi were harvested 4d. post-SE and were subjected to either CUT&RUN-seq or RNA-seq. **(B)** Volcano plot of differential enrichment analysis of H3K27me3 peaks between naive and 4d. post-SE hippocampi (n=3442 differential peaks; FDR < 0.05). Blue and red circles represent peaks enriched and depleted 4d. post-SE, respectively. **(C)** Normalized heatmap and metaplot (RPKM) of average H3K27me3 signal in naive and 4d. post-SE hippocampi across differentially enriched and depleted peaks. Plots are centered on the peak summit and extend ± 3kb. Metaplots represent average normalized signal across all peaks in differentially enriched (blue line) and depleted (red line) peaks per condition. **(D)** Venn diagram showing overlap of H3K27me3 peaks enriched 4d. post-SE with H3K27me3 peaks in the naive hippocampus. Indigo represents pre-marked loci and purple represents de novo loci. **(E)** Scatter plot comparing naive (x-axis) and 4d. post-SE (y-axis) H3K27me3 signal for all quantified H3K27me3 peaks. **(F)** Scatter plot highlighting pre-marked and de novo H3K27me3 enriched peaks.

We sought to determine whether H3K27me3 was being added to loci pre-marked with methylation in naive brains or whether H3K27me3 was being deposited at new loci (de novo sites). To test this, we analyzed the genomic overlap of all enriched H3K27me3 peaks with the consensus set of H3K27me3 peaks in the naive hippocampus. We find that enriched H3K27me3 peaks were nearly equally distributed between loci pre-marked by H3K27me3 in the naive hippocampus (n=1335) and loci that were de novo methylated (n=1324) (Fig. 1D, S2 Data), suggesting that H3K27me3 deposition induced by SE did not require existing H3K27me3. Plotting H3K27me3 signal in naive and post-SE hippocampi shows that de novo loci tend to exhibit greater magnitude of increase in H3K27me3 signal compared to pre-marked loci (Fig. 1E-F). Taken together, these suggest that SE induces H3K27me3 enrichment at both pre-marked and de novo loci.

### H3K27me3 induction is associated with repression of network excitability genes

To determine the effect of H3K27me3 induction post-SE on transcription, we integrated H3K27me3 CUT&RUN data with bulk RNA-seq data from naive and post-SE hippocampi collected in tandem with the epigenomic dataset. Differential expression analysis identified 6225 differentially expressed genes (p-adj. < 0.05, fold-change (FC) cut-off = 1.2), of which 3482 were induced and 2743 were repressed post-SE (Fig. 2A, S3-4 Data). To compare H3K27me3 enrichment with RNA expression, we first annotated all enriched H3K27me3 peaks to genes based on their overlap with closest genomic feature (see methods, S5 Data). Most H3K27me3 enriched peaks could be annotated to either gene promoters (73.9%) or gene bodies (15.7%) (Fig. 2B). In total, H3K27me3 enriched peaks represented 2022 unique genes of which 1130 were pre-marked by H3K27me3 while 892 were de novo H3K27 tri-methylated (Fig. 2C). We then categorized genes into those that were induced, repressed, showed no change, or were undetectable and compared the distributions of these categories between pre-marked and de novo annotated genes (Fig. 2C-E). Approximately 50% of genes annotated to H3K27me3 enriched peaks showed no change in RNA expression between naive and 4d. post-SE hippocampi and were not preferentially associated with either pre-marked or de novo loci. In contrast, 287 genes representing 14% of H3K27me3 enriched genes were undetected by RNA-seq, likely corresponding to silenced genes. These undetected genes were significantly associated with pre-marked loci and exhibit the highest levels of H3K27me3 deposition across the promoter and gene body when compared to genes in other categories (Fig. 2E). Finally, 519 genes (25% of H3K27me3 enriched genes) were repressed post-SE. Repressed genes exhibited less H3K27me3 across the gene body compared to undetected genes and genes with no change in RNA and were significantly associated with de novo H3K27 methylation. Finally, a subset of genes were induced post-SE (n=168; 8% of H3K27me3 enriched genes) and these were not preferentially associated with either pre-marked or de novo loci. H3K27me3 signal and RNA tracks comparing naive and 4d. post-SE hippocampi for representative genes for each of the 4 RNA expression categories are shown in Fig. 2F-I. Taken together, these findings suggest that H3K27me3 deposition post-SE may function to repress a subset of genes through predominantly de novo methylation but, H3K27me3 deposition post-SE overall, does not necessarily lead to gene repression.

**Fig. 2.**
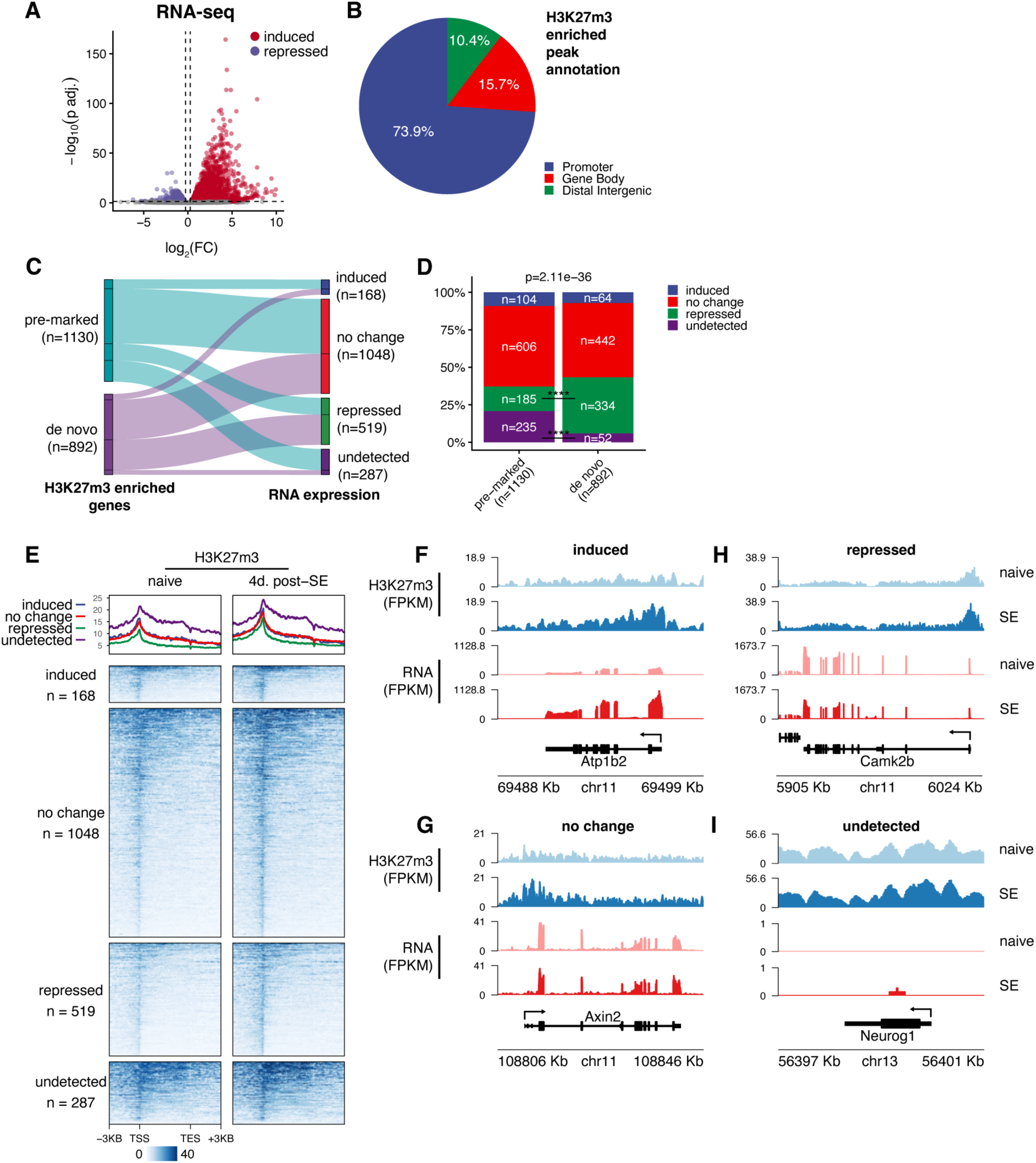
Integration of CUT&RUN and RNA-seq reveals complex associations of H3K27me3 and expression. **(A)** Volcano plot showing differential gene expression analysis between naive and 4d. post-SE hippocampi (n=6225 differentially expressed genes; p adj. < 0.05; fold-change (FC) threshold = 1.2). Blue and red circles represent genes induced and repressed 4d. post-SE, respectively. **(B)** Genomic distribution of H3K27me3 peaks enriched 4d. post-SE. Promoters were defined as within 3kb upstream to 200bp downstream of the gene transcription start site (TSS). **(C-D)** Sankey diagram summarizing distribution of RNA expression patterns of pre-marked or de novo H3K27me3 enriched genes. Distributions are significantly different between genes annotated to pre-marked and de novo H3K27me3 enriched peaks (X^2^ test, p=2.11e-36). Post-hoc test shows that repressed and undetected genes are asymmetrically distributed between pre-marked and de novo loci (****; p < 0.0001). **(E)** Normalized heatmap and metaplot (RPKM) of average H3K27me3 signal in naive and 4d. post-SE hippocampi across genes annotated to H3K27me3 enriched peaks post-SE, clustered by RNA expression pattern. Signal is plotted over genes scaled to 5kb from transcription start site (TSS) to transcription end site (TES). Plots are extended -3kb from TSS and +3kb from TES. Metaplots represent the average signal across all plotted regions for each RNA expression cluster. **(F-I)** Average H3K27me3 CUT&RUN (FPKM) and RNA coverage (RPKM) in naive and 4d. post-SE hippocampi for representative genes in each RNA expression cluster (E: induced, F: no change, G: repressed, H: undetected). Arrows indicate direction of gene transcription.

To gain insight into the cellular pathways and processes represented by H3K27me3 target genes, we performed ontology analysis against the KEGG [20], Reactome [21], Biocarta [22,23], and Hallmark [24] databases (S6-10 Data). Ontology results from each database were then combined and clustered based on overlap of genes for each annotation term to summarize ontology analyses results across different databases [13]. We identified 5 clusters of gene ontology terms for H3K27me3 enriched genes with no change in expression (Fig. S1A, S6 Data). Clusters 1 (Basal Cell Carcinoma) and 3 (Constitutive Signaling by Aberrant PI3K in Cancer) comprised the majority of terms annotated to this gene set and were related to developmental signaling programs that are commonly dysregulated in cancer such as Wnt [25,26] and Hedgehog signaling [27,28] (Fig. S1A, S6 Data). The remaining clusters were related to various physiological processes including Axon Guidance (Cluster 2), Chondroitin Sulfate Biosynthesis (Cluster 4), and Cytochrome P450 Arranged by Substrate Type) (Fig. S1A, S6 Data). We also examined genes induced post-SE that were either H3K27me3 enriched or unmethylated (Fig. S1B-C, S7-8 Data). Induced genes that were also H3K27me3 enriched yielded ontological terms related to immunosuppressive functions typically described in cancer pathogenesis including TGF-beta signaling and epithelial-to-mesenchymal transition (Fig. S1B, S7 Data) [29]. Unmethylated induced genes yielded ontological clusters related to immune signaling including innate immune system, interferon gamma response, and adaptive immune system as well as various cancer-related pathways such as extracellular matrix organization, G2M checkpoint, and epithelial-to-mesenchymal transition (Fig. S1C, S8 Data).

In contrast, we identified 8 clusters of gene ontology terms with broadly neuronal related functions for H3K27me3 enriched, RNA repressed genes (Fig. 3A, S9 Data). The top scoring cluster (Neuronal System) represented a broad functional category characterized by genes typically expressed in neuronal cells and systems. Clusters 2 (Transmission Across Chemical Synapses), 4 (Axon Guidance), and 7 (Phase 0 Rapid Depolarisation) represented genes generally annotated to synaptic signal transmission such as various ion channels and neuroactive ligand receptors (Fig. 3A, S9 Data). We also find clusters corresponding to other neuronal processes such as axon guidance and adherens junctions interactions. Interestingly, cluster 3 (Regulation of Insulin Secretion) comprises terms annotated to insulin secretion, but importantly, these terms are annotated to genes involved in the same vesicle release pathways in neurotransmitter release at the pre-synaptic zone [30,31] (S9 Data). We further analyzed predicted protein-interaction networks from the set of H3K27me3 target neuronal signaling genes we identified from ontology analysis using the STRING database [32] (Fig. 3B). H3K27me3 target neuronal signaling genes were enriched for interaction networks corresponding to various proteins regulating neurotransmission including Ca2^+^ and K^+^ channels, N-methyl-D-aspartate (NMDA) receptors, and Ca^2+^/Calmodulin-dependent Kinases (CAMKs) (Fig. 3B). We also analyzed repressed genes that did not exhibit increased H3K27me3 deposition and similarly identified clusters of terms broadly related to neuronal functions including vesicle-mediated transport, protein-protein interaction at synapses, and long-term depression and potentiation (Fig. S1D, S10 Data). Taken together, transcriptionally repressed genes, of which a subset are direct H3K27me3 targets, represent pathways which may affect neuronal signaling and network excitability.

**Fig. 3.**
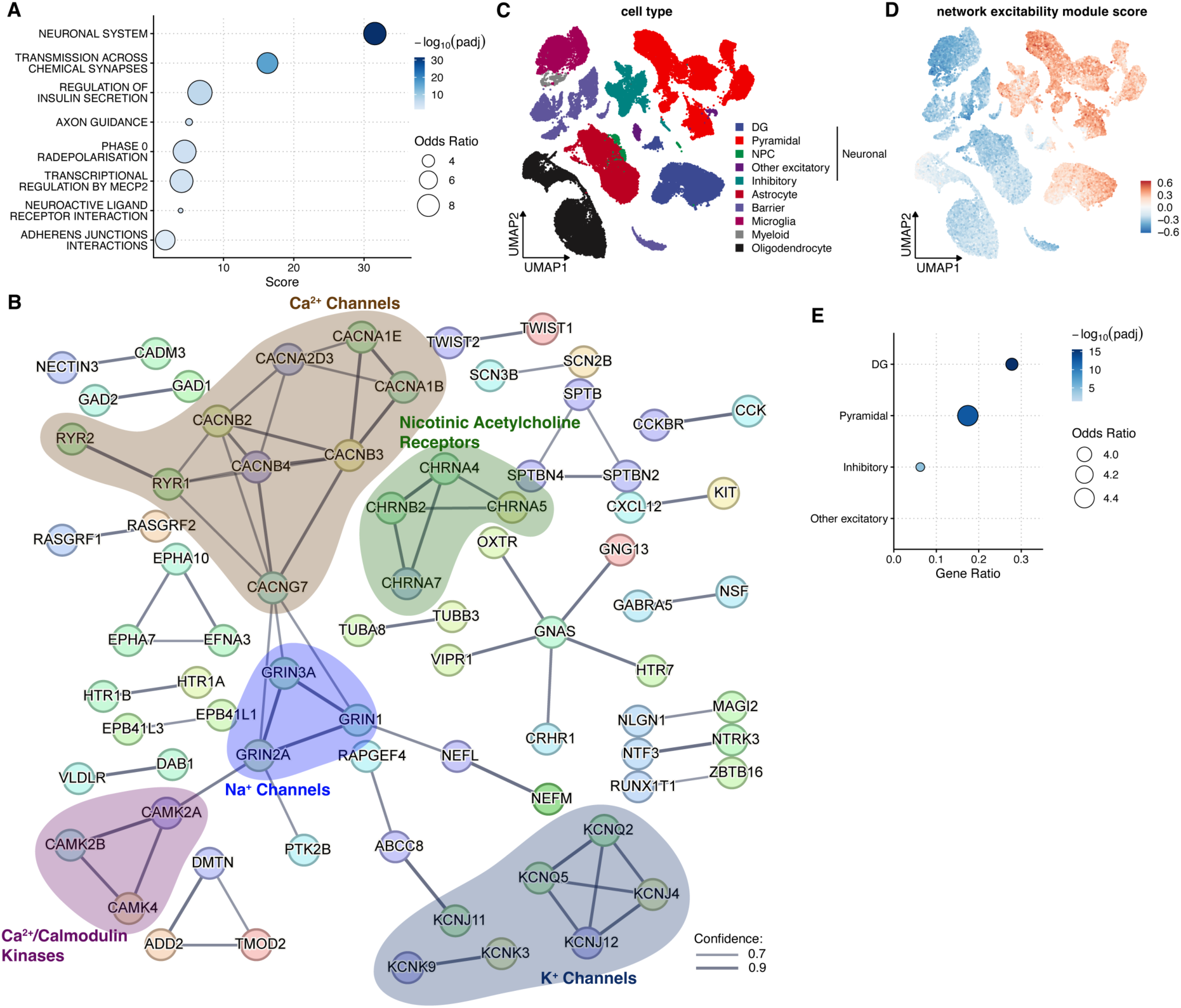
Repressed genes associated with H3K27me3 deposition represent pathways related to network excitability. **(A)** Summarized ontology analysis of H3K27me3 enriched RNA repressed genes reveals enrichment for pathways involved in neuronal signaling and network function. Ontology analysis was summarized as described in methods. Color is scaled to the -log_10_(padj) of the top-scoring term per cluster. Size is scaled to the Odds Ratio. **(B)** Physical interaction network of target neuronal signaling/network excitability genes identified from (A). Network was trimmed to exclude interactions with Confidence < 0.7 and disconnected nodes in the network. Local clusters of various ion channels and Ca^2+^/Calmodulin-dependent kinases are highlighted. **(C-D)** Meta-analysis of snRNA-seq data from a pilocarpine model of SE[13] shows that target genes identified from ontology analysis are enriched in neuronal cells. **(E)** Down-regulated genes in DG, Pyramidal, and Inhibitory cells are enriched for the H3K27me3 enriched RNA repressed genes related to network excitability and neuronal signaling.

We hypothesized that the functional consequences of H3K27me3 deposition would likely be mediated by target genes whose expression was altered post-SE. Thus, we focused our analysis on H3K27me3 enriched, RNA repressed genes, as these would be consistent with the canonical repressive function of H3K27me3. Given that bulk CUT&RUN-seq and RNA-seq provide aggregate data across cell types, we sought to determine the cell-type specific expression of H3K27me3-enriched, RNA-repressed genes we identified from ontology analyses. We analyzed previously generated single nuclei RNA-seq (snRNA-seq) data from naive and 4d. post SE mouse hippocampi [13]. Using this atlas of single cell molecular changes post SE, we calculated the relative expression of the set of 241 network excitability genes across all cells using the scanpy [33] implementation of the expression scoring method described by Tirosh et al [34]. Analysis of gene score across all cells shows that target network excitability genes are broadly enriched in terminally differentiated neuronal cells compared to non-neuronal cells and neural progenitor cells (NPCs) (Fig. 3C-D, Fig. S2). Thus, we focused further analyses on neurons. We then sought to determine which neuronal cell types exhibited SE-dependent down-regulation of the network excitability genes we identified from our analysis of bulk data. To this end, we identified significantly repressed genes for each neuronal cell type from pseudobulked expression data (see methods) and compared the overlap with the set of 241 target network excitability genes (Fig. 3E). Of these, dentate granule and pyramidal cells showed the most significant degree of overlap with the target network excitability genes.

Taken together, these suggest that H3K27me3 deposition post-SE may function in part to repress genes involved in regulating neuronal function and signaling in excitatory neurons in the hippocampus, thereby modulating network excitability acutely after injury.

### EZH2 inhibition post-SE increases network excitability acutely

To test whether inhibition of EZH2 methyltransferase function post-SE would alter the SE-dependent repression of target genes controlling network function, we treated both naive mice and mice induced to SE with the EZH2 inhibitor UNC1999 [12,35]. Mice were injected with UNC1999 or vehicle control 6, 24, and 48 hours post-SE resulting in the following 4 conditions: naive + vehicle, SE + vehicle, naive + UNC1999, SE + UNC1999 as in Khan et al. [12] (Fig. 4A). H3K27me3 was enriched at the TSS of target genes in SE + vehicle samples compared to naive + vehicle controls, as expected (Fig. 4B). We also observed diminution of H3K27me3 in SE + UNC1999 compared to SE + vehicle samples (p = 0.0253, 1.2 Fold reduction), suggesting that UNC1999 treatment resulted in attenuation of H3K27me3 enrichment post-SE (Fig. 4B-C). Similarly, analysis of RNA expression shows that network excitability genes are repressed in SE + vehicle samples compared to naive + vehicle, and SE + UNC1999 samples exhibit diminished reduction in RNA expression compared to SE + vehicle (p = 2.14e-77) (Fig. 4C-D).

**Fig. 4.**
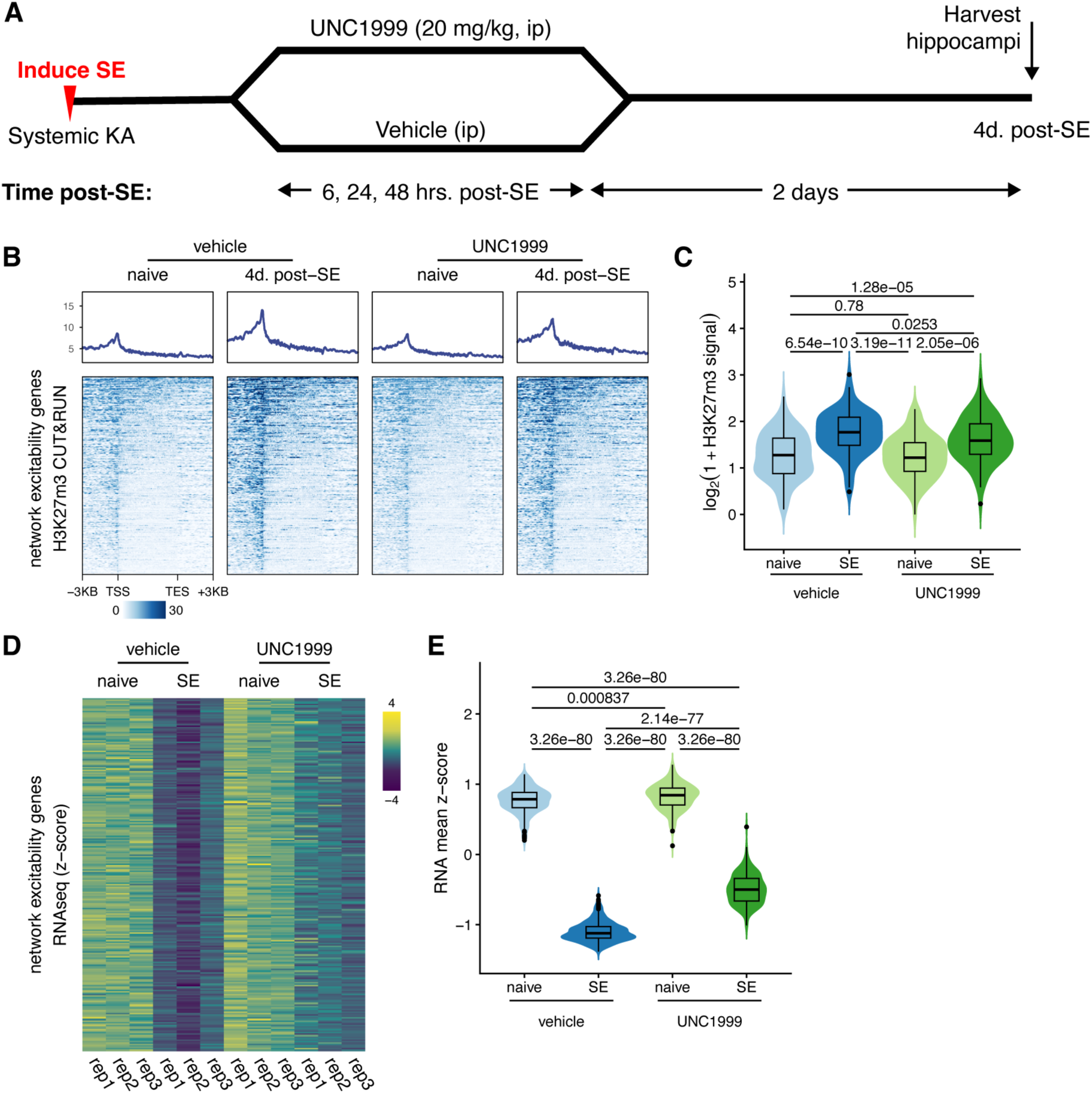
UNC1999 treatment attenuates SE-dependent H3K27me3 deposition and RNA repression of network excitability genes. **(A)** Animals were induced to SE by systemic KA administration and treated with either vehicle or UNC1999 6, 24, and 48 hrs. post-SE. Hippocampi were harvested 4d. post-SE for genomic and transcriptomic analyses. **(B)** Normalized heatmap and metaplot (RPKM) comparing average H3K27me3 signal of network excitability genes of interest in naive and 4d. post-SE hippocampi treated with either vehicle or UNC1999. **(C)** Violin plot comparing distributions of mean H3K27me3 signal at TSS ± 200bp of network excitability genes. UNC1999 treated animals show marginally decreased H3K27me3 post-SE compared to vehicle treated animals. Values were subjected to log_2_(1 + x) transformation for plotting.**(D)** Normalized heatmap comparing RNA expression of target network excitability genes in naive and 4d. post-SE hippocampi treated with either vehicle or UNC1999. Expression was z-score normalized per gene. **(E)** Violin plot comparing distributions of mean z-score normalized RNA expression per condition. Target genes exhibit attenuated repression post-SE in UNC1999 treated animals compared to vehicle control. For comparisons of H3K27me3 signal and RNA expression between different conditions, Kruskal-Wallis rank sum test was performed followed by Wilcoxon rank sum test with continuity correction for pairwise comparisons. Benjamini-Hochberg Procedure was used for multiple comparisons correction and adjusted p-values are displayed in plots.

Given that UNC1999 treatment attenuated the repression of genes predicted to control neuronal signaling and network function, we hypothesized that UNC1999 treatment would also functionally alter network excitability post-SE. To test this, we measured network excitability via seizure threshold, measured as the time to generalized tonic-clonic seizure (GTCS) upon exposure to flurothyl [12,13] (Fig. 5A). Vehicle-treated mice showed no change in time to GTCS between naive baseline and 4d. post-SE measurements (Fig. 5B), suggesting that under normal conditions, mice that undergo SE maintain their baseline network excitability acutely after injury. In contrast, mice treated with UNC1999 show a statistically significant reduction in time to GTCS at 4d. post-SE compared to naive baseline (Fig. 5B). To rule out the possibility that UNC1999 treatment reduces seizure threshold independent of SE, we measured baseline and post-treatment seizure threshold in healthy mice (ie. mice that did not receive kainate to induce SE but given saline instead) treated with UNC1999 (Fig. S3) and found no change in time to GTCS after UNC1999 treatment. Taken together, these suggest that UNC1999 treatment post-SE abrogates the maintenance of network excitability acutely after injury.

**Fig. 5.**
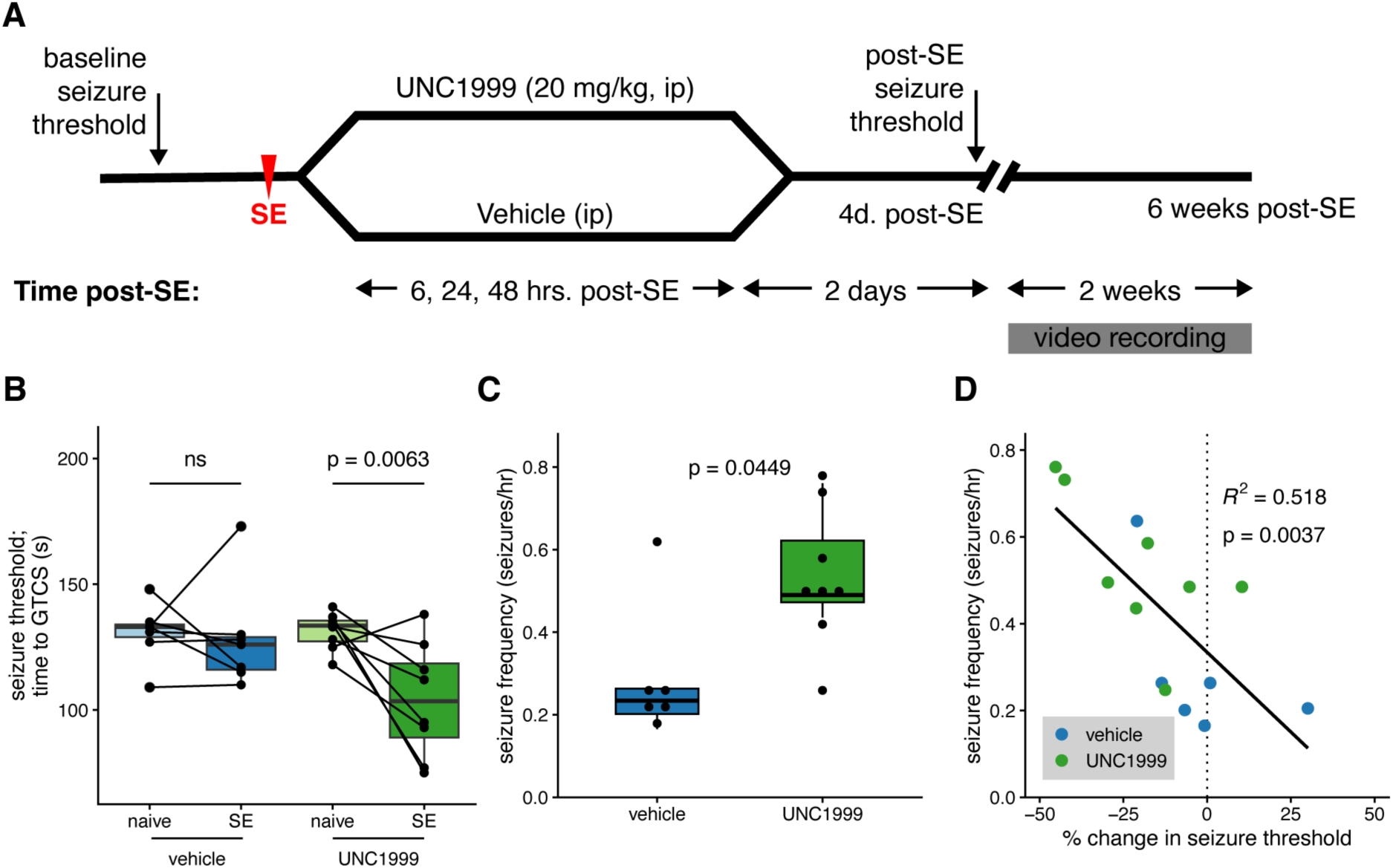
UNC1999 treatment post-SE increases network excitability acutely and disease severity chronically. **(A)** Animals were treated with either vehicle or UNC1999 as in Fig.4A. For each mouse, seizure threshold was recorded prior to SE to determine naive baseline threshold and again 4d. post-SE. Behavioral seizures were monitored by video for 2 weeks from weeks 5-6 post-SE. **(B)** Seizure threshold (time to GTCS) is significantly decreased 4d. post-SE only in animals treated with UNC1999 but not in vehicle treated animals (2-way ANOVA with repeated measures followed by Tukey’s multiple comparisons test). Lines connecting points indicate repeated measures for an individual mouse. **(C)** Seizure frequency during the chronic phase was significantly higher in animals treated with UNC1999 compared to vehicle control (Wilcoxon rank sum test). **(D)** Percent change in seizure threshold (4d. post-SE – naive) is negatively correlated with seizure frequency in the chronic period (Pearson correlation). Points indicate individual mice.

We then sought to determine how this observed change in network excitability acutely after injury correlated with disease severity in the chronic period. To this end, we monitored behavioral seizures by video recording in the same cohort of vehicle and UNC1999 treated mice between 5-6 weeks post-SE. Mice treated with UNC1999 showed higher seizure frequency compared to vehicle treated mice (Fig. 5C), corroborating our previous findings [12]. Moreover, change in seizure threshold from naive baseline to 4d. post-SE was significantly negatively correlated with seizure frequency (Fig. 5D) such that an acute reduction in seizure threshold was associated with increased spontaneous seizure frequency in the chronic period. This suggests that maintenance of network excitability acutely after injury may affect disease severity chronically. In summary, EZH2 inhibition via UNC1999 treatment post-SE attenuated both H3K27me3 deposition and RNA repression of genes controlling network excitability. Functionally, EZH2 inhibition disrupted the maintenance of network excitability acutely which was correlated with exacerbated disease severity chronically.

## DISCUSSION

Herein we describe a putative mechanism by which EZH2 mediates control of network excitability acutely after epileptogenic injury. We show that H3K27me3 deposition acutely post-SE is associated with transcriptional repression of genes encoding proteins regulating various aspects of neuronal signaling and network function in the mouse hippocampus. Inhibition of EZH2 H3K27 methylase activity attenuates H3K27me3 enrichment at these genes and diminishes their repression post-SE. Functionally, EZH2 inhibition acutely post-SE increases network excitability and this is associated with exacerbated disease severity chronically. Taken together, these suggest that H3K27me3 deposition post-SE may function in part to maintain network excitability after injury, thereby lowering seizure probability chronically.

Acutely post-SE, H3K27me3 is enriched both at loci with pre-existing H3K27me3 and de novo loci (Fig. 1). It is well-known that H3K27me3 can propagate through a read-and-write mechanism where an initial H3K27me3 locus recruits PRC2 and stimulates further H3K27me3 deposition at neighboring nucleosomes both in *cis* and *trans*, resulting in spreading of H3K27me3 [36–38]. It is possible that upon SE-dependent EZH2 induction, PRC2 is recruited to a subset of pre-marked H3K27me3 loci in the hippocampus resulting in further enrichment of H3K27me3 at these loci. In contrast, the mechanisms for de novo H3K27me3 deposition are less understood and remain controversial [39–42]. Earlier work in mouse embryonic stem cells suggests that SUZ12 is sufficient to bind unmethylated loci to recruit and stabilize PRC2 assembly, thereby leading to de novo H3K27 methylation [39]. A similar mechanism may occur in the post-SE hippocampus where SUZ12 facilitates PRC2 recruitment and activity at de novo targets upon SE, but this needs to be experimentally validated. Others have also shown that H2AK119 mono-ubiquitylation via PRC1 is necessary for recruitment of PRC2 and subsequent de novo establishment of H3K27 methylation [40]. However, we could not find evidence of changes in H2AK119ub upon SE (S4 Fig) and whether H2AK119 mono-ubiquitylation contributes to SE-dependent de novo H3K27me3 deposition is unknown. Finally, it is important to note that the current understanding of de novo H3K27me3 deposition is derived almost exclusively from studies of H3K27me3 establishment and maintenance during cell differentiation and there may be other distinct mechanisms for de novo H3K27 methylation in terminally differentiated cells.

It should be noted that this study lacks direct profiling of chromatin occupancy by EZH2 and other PRC2 sub-units. Mapping genome-wide occupancy of EZH2 in naive and post-SE hippocampi would provide direct evidence for identification of EZH2 target genes. However, we were unable to profile EZH2 nor SUZ12 chromatin occupancy in post-SE whole hippocampi using CUT&RUN-seq. Although we were able to consistently identify EZH2 and SUZ12 CUT&RUN peaks in naive mouse hippocampi, CUT&RUN against the same epitopes with the same antibodies in post-SE hippocampi yielded CUT&RUN signal that was indistinguishable from background IgG (Fig. S5A). This trend was consistent even when we profiled the transcription factor CTCF (Fig. S5B) suggesting that post-SE hippocampi were not permissive to transcription factor/co-factor CUT&RUN-seq. We speculate that extensive DNA damage and cell death induced by SE [43] generates excessive background signal from free DNA fragments that overwhelms the sparse signal from chromatin-bound TFs and co-factors but is not sufficient to dilute the signal from more abundant targets like histone modifications.

Instead, we profiled H3K27me3 as a proxy for EZH2 activity, as H3K27 methylation is the enzymatic output of EZH2. The only known H3K27 methylases in mammals are the homologous proteins EZH1 and EZH2 [44,45], and only EZH2 is induced acutely post-SE [12]. Moreover, we do not observe changes in expression of known H3K27 demethylases (KDM6A and KDM6B) [46–49] (S3-4 Data), suggesting that alterations in H3K27me3 we observe are not due to differential demethylase activity. Thus, it is likely that alterations in H3K27me3 that we measured correspond to increased EZH2 activity.

Our analysis of H3K27me3 deposition in tandem with RNA expression reveal complex associations between H3K27me3 and gene expression. We find that over 77% of genes with increased H3K27me3 post-SE showed either no change in RNA expression or were undetected by RNA-seq (Fig. 2). Given the canonical function of H3K27me3 in maintenance of gene silencing, it is tempting to speculate that SE-dependent H3K27me3 deposition at these genes is important for preventing ectopic expression post-injury. However, EZH2 inhibition post-SE did not appear to alter the expression of most of these genes (∼90%) (S14 Data) as measured by RNA-seq suggesting that this is not the case. It is important to note though, that most of these genes are expressed in low to undetectable quantities in the naive hippocampus, making precise quantification through typical RNA-seq workflows difficult (S3-4 Data, S14 Data) [50]. Whether SE-dependent H3K27me3 deposition at these genes is functional or is simply a by-product of EZH2 induction is still unknown. In contrast, we identified over 500 genes in which H3K27me3 deposition was associated with concomitant transcriptional repression (Fig. 2) and we focused subsequent analysis here. Among these, we identify a module of genes annotated to functions involved in modulation of neuronal signaling. These included genes underlying excitatory neurotransmission that have been extensively studied for their contributions to seizure generation and various epilepsies such as N-methyl-D-aspartate (NMDA) receptors [51], nicotinic acetylcholine receptors (nACHRs) [52], voltage-gated Na^+^ channels [53], voltage-gated Ca^2+^ channels [54], and calcium/calmodulin-dependent kinases (CAMKs) [55,56]. Also included were genes encoding molecules regulating axon guidance and synaptic organization which are critical processes involved in epilepsy pathogenesis [57,58]. EZH2 inhibition post-SE attenuated H3K27me3 deposition and diminished transcriptional repression of this network excitability gene module (Fig. 4) and this epigenomic and transcriptomic alteration was associated with increased network excitability post-SE (Fig. 5). In contrast, mice that had undergone SE but were treated with vehicle instead of UNC1999 showed no statistically significant change in network excitability. These suggest that under normal conditions, the hippocampus maintains network excitability acutely after injury and this maintenance requires EZH2 methyltransferase activity.

Changes in network excitability acutely after injury correlated with disease severity weeks later in the chronic period. Specifically, we show that increased network excitability post-SE brought about by EZH2 inhibition is associated with exacerbated seizure frequency during the chronic period. We hypothesize that EZH2 induction mediates a homeostatic mechanism to maintain either intrinsic neuronal excitability or overall network excitability in the hippocampus in response to SE. This is eventually overcome by epileptogenic alterations in the brain that ultimately leads to spontaneous seizures (i.e. epilepsy). However, EZH2 inhibition prevents this endogenous maintenance of baseline excitability, effectively accelerating disease progression by lowering the barrier/threshold required for epileptogenic alterations to overcome to generate spontaneous seizures.

Work from our lab and others identified EZH2 as a critical factor regulating gene expression acutely post-SE with a net protective effect against disease though the exact mechanism of function was unknown [12,13]. We previously proposed a model in which EZH2 functioned to temper the induction of inflammation in the hippocampus acutely post-SE via repression of JAK/STAT signaling components [13]. Here, we integrate epigenomic and transcriptomic profiling to interrogate EZH2 mechanism of function post-SE. Despite a robust elevation of JAK/STAT signaling in the absence of EZH2 post SE compared to WT animals our analysis herein failed to find deposition of H3K27me3 at any of the JAK nor STAT protein encoding loci (S5 Data). This suggests that EZH2 is either i) indirectly controlling JAK/STAT signaling or ii) directly controlling JAK/STAT signaling via regulation of hitherto unknown components of the pathway.

One of the major limitations of this study is our use of bulk methods for epigenomic and transcriptomic profiling of whole hippocampal tissue. The hippocampus comprises a heterogenous population of cells with cell-type specific as well as spatially-distinct transcriptomes and functions [59–61]. Because we did not enrich for specific cell populations, the epigenomic and transcriptomic alterations brought about by SE that we observe represent aggregate data from all hippocampal cell types. One consequence of this is that the transcriptomic contribution of cell types that represent smaller populations in the overall tissue, but are nonetheless important in disease pathology, will be masked by more abundant cell types. For example, inhibitory interneurons represent only roughly 10-15% of neuronal cells in the hippocampus [62]. Despite this, interneuron loss or dysfunction mediates profound changes in the balance of excitation and inhibition in the brain and is associated with seizure generation and epileptogenesis [63–65]. In such cases, cell-type specific analyses or single-cell/nuclei sequencing methods would resolve the specific contribution of each cell type to the whole population. To overcome these limitations, we supplement our bulk epigenomic and transcriptomic data by integrating analysis of a previously generated mouse pilocarpine SE snRNA-seq dataset [13]. Specifically, we show that the module of neuronal signaling and network excitability target genes we identify from bulk data are enriched in neuronal cells. Moreover, analysis of expression pseudobulked per cell type shows that genes repressed in dentate granule and pyramidal cells post-SE exhibit significant overlap with the neuronal signaling and network excitability gene module. Taken together with our functional results, these support the hypothesis that EZH2 controls neuronal signaling primarily in excitatory neurons to maintain network excitability acutely after SE.

Finally, the study only profiles H3K27me3 and RNA expression at an early stage of disease progression (4d. post-SE). Whether the changes in H3K27me3 deposition or RNA expression we observe at 4d. post-SE persist throughout the acute phase or into the chronic phase are unknown. Work to define cell-type specific epigenomic and transcriptomic trajectories throughout epileptogenesis is currently ongoing.

In summary, we identify a putative mechanism underlying the protective effect of EZH2 induction post-SE by using an integrative multi-modal approach combining epigenomic, transcriptomic, and functional analyses. We show through orthogonal lines of evidence that EZH2 induction may function to maintain network excitability to lower seizure probability chronically. Further work to decipher the specific circuits and pathways that mediate EZH2 control of network excitability acutely post-SE may provide potential targets for disease modifying treatment.

## METHODS

### Animal care

All animal procedures were performed with approval from the University of Wisconsin-Madison School of Medicine and Public Health Institutional Animal Care and Use Committee and according to NIH national guidelines and policies.

#### Repeated low-dose KA model of status epilepticus

Mice were weighed and singly housed in polycarbonate observation chambers for the duration of SE induction and observation. Mice were injected with synthetic kainic acid (KA) (#7065, Tocris Bioscience, Bristol, United Kingdom) dissolved in 0.9% saline to induce SE. Mice were injected intraperitoneally (i.p.) at twenty-minute intervals at 7.5mg/kg for the first three injections and 5.0mg/kg for subsequent injections until reaching SE. An injection was skipped if an animal experienced two or more Class V or VI seizures within a single twenty-minute interval. Injections resumed the next interval unless the animal reached SE. Throughout SE induction, behavioral seizures were observed and recorded based on a modified Racine Scale [66] as in the table below:

**Table 1.**
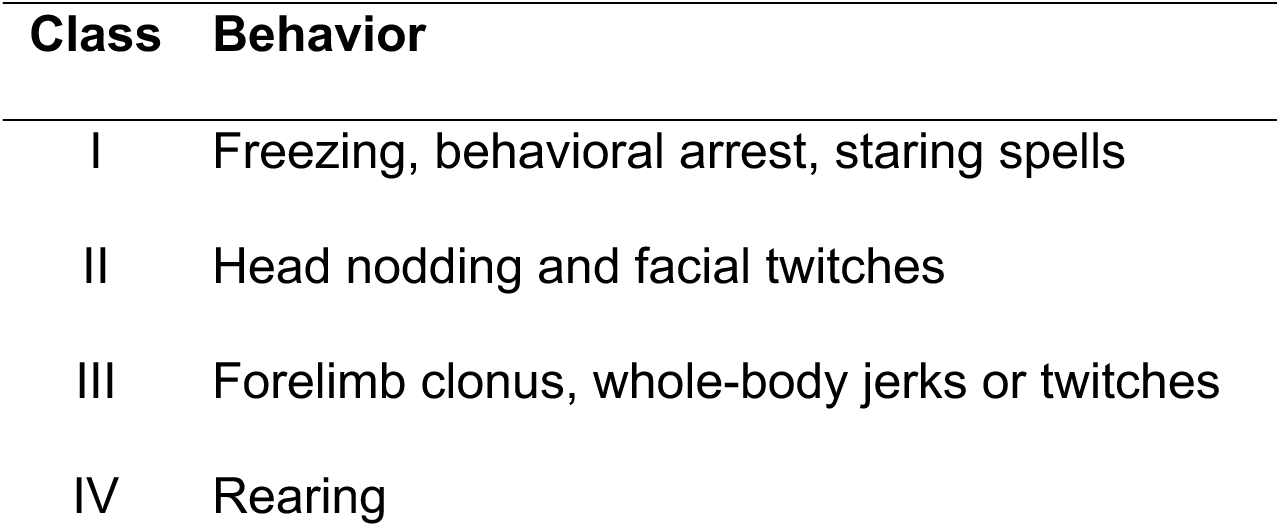
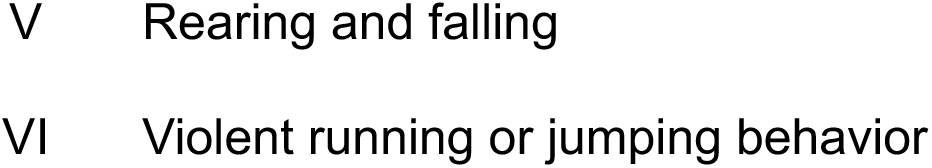
Modified racine scale.

Animals were considered to have reached SE after occurrence of at least five Class V or VI seizures within a 90-minute window. KA mice were monitored for 1-2 hours after SE was achieved. After SE induction, animals were returned to home cages and provided with soft gel food for recovery. In the days following injection, animals were weighed and injected subcutaneously (s.c.) with 0.9% saline if body weight had decreased by more than 10% of initial body weight prior to SE induction.

#### UNC1999 treatment

UNC1999 solution (2.5mg/mL) (#14621 Cayman Chemical, Ann Arbor, MI) was diluted in 4% N,N-Dimethylacetamide (DMA; Sigma Aldrich), 5% solutol (Sigma Aldrich), and 0.9% saline as previously described [12]. Mice were treated with 20mg/kg UNC1999 or vehicle control at 6, 24, and 48 hrs. post-SE. For SFig. 3, mice were treated with UNC1999 at 6, 24, and 48 hrs. post-sham SE induction.

#### Seizure threshold testing

Flurothyl seizure threshold tests were performed on mice using 100% bis(2,2,2- trifluoroethyl) ether (#287571, Sigma-Aldrich, St. Louis, MO, USA). All tests were performed in a fume hood. Mice were placed in an airtight 10L Plexiglass chamber (8.5”x10.75”x6.75”). Flurothyl was infused into the chamber via a peristaltic pump (40uL/minute) onto a piece of Whatman #1 filter paper placed at the top of the chamber. Flurothyl infusion was continued until mice were induced to generalized tonic clonic seizure (GTCS) observed as a complete loss of postural/motor control. Seizure threshold was measured as the time to reach GTCS from the start of flurothyl infusion. After GTCS, mice were rapidly removed from the testing chamber and transferred to a recovery cage until normal behavior resumed (approximately 10 min). Mice were returned to home cages after recovery. All seizure threshold tests were video recorded and seizure behavior was scored blinded observer. For Fig. 5, two flurothyl seizure threshold tests were performed per mouse; baseline seizure threshold was recorded one week prior to kainate injections and post-SE seizure threshold was recorded 4 days post-SE. For S1 Fig., two flurothyl seizure threshold tests were performed per mouse; baseline seizure threshold was recorded 1 week prior to sham SE induction and UNC1999 treatment, and post-sham seizure threshold was recorded 4 days after sham SE induction. Seizure threshold tests were performed at approximately the same time of day (10 AM) and mice were returned to home cages before 5 PM.

#### Tissue isolation and storage

For hippocampal isolation for molecular experiments, animals were sacrificed by decapitation. Whole hippocampal hemispheres were dissected and flash-frozen in dry ice and stored at -80°C.

#### Behavioral seizure monitoring

Post-SE mice were monitored for behavioral seizures by video recording from 5- 6 weeks post-SE. Mice were singly housed in observation chambers during recording sessions with access to food and water. Two recording sessions of 5-6 hrs. duration were done per week for a total of 4 recording sessions. Behavioral seizures were scored by blinded observers. Seizure frequency for each mouse was calculated as the number of observed seizures across all recording sessions divided by the total duration of recording.

### H3K27me3 CUT&RUN

#### CUT&RUN-sequencing

Flash frozen hippocampi were thawed for 30 seconds at 37°C and washed twice in room temperature 1x PBS supplemented with EDTA-free protease inhibitor cocktail. Hippocampi were dissociated by trituration with a wide-bore P1000 tip (10 draws) followed by a wide-bore P200 tip (10 draws) in 0.5mL 1X PBS with protease inhibitor cocktail. Tissue suspension was further dissociated in 1mL dounce with the loose pestle (10 strokes). Cells were lightly fixed in 0.1% formaldehyde for 2 min. at RT. Cross- linking was quenched by addition of 1.25M glycine to final concentration of 55mM for 5 min. at RT. Cross-linked cells were pelleted by centrifugation at 600xg for 5 min. at 4°C in swinging bucket rotor. Cross-linked cells were washed twice with 1x PBS + PIC and pelleted with centrifugation as above. Nuclei were isolated by hypotonic lysis in 2.5mL NE1 (20mM HEPES-KOH pH 7.9, 10mM KCl, 0.5mM spermidine, 0.1% Triton X-100, 20% glycerol, 1x Roche cOmplete EDTA-free protease inhibitor) per hippocampus for 10 min on ice. Nuclei were pelleted by centrifugation at 1300xg for 8 min. at 4°C in swinging bucket rotor and resuspended in room-temperature CUT&RUN Wash150 buffer (20mM HEPES pH 7.5, 150mM NaCl, 0.5mM spermidine, 1x Roche cOmplete EDTA-free protease inhibitor) at 1×10^6^ nuclei/mL. For nuclei/cell counting, nuclei were stained with 0.1% Trypan Blue and counted on a standard hemacytometer. For each CUT&RUN reaction, 100,000-250,000 nuclei were bound to 10uL of ConA paramagnetic beads (#21-1411, EpiCypher, Durham, NC, USA) in 0.2mL PCR strip-tubes and incubated on a nutator at RT for 10 min. For subsequent wash and resuspension steps, separation of bead-bound nuclei was achieved by incubation of beads on magnetic separation rack for 1-2 min. After bead-binding, nuclei:bead suspension was resuspended in 100 uL Antibody150 buffer (20mM HEPES pH 7.5, 150mM NaCl, 2mM EDTA, 0.5mM spermidine, 0.01% Digitonin, 1x Roche cOmplete EDTA-free protease inhibitor) and incubated with either anti-H3K27me3 (1:50) (#9733, Cell Signaling Technology, Inc., Danvers, MA, USA) or Rabbit IgG isotype control (1:20) (#66362, Cell Signaling Technology, Inc., Danvers, MA, USA) overnight on nutator at 4°C. Nuclei were washed twice in 200uL Dig150 Buffer (20mM HEPES pH 7.5, 150mM NaCl, 0.5mM spermidine, 0.01% Digitonin, 1x Roche cOmplete EDTA-free protease inhibitor) to remove excess antibody. Nuclei were resuspended in 100uL Dig150 Buffer + pA/G- MNase (700ng/mL) pre-mix and incubated on nutator for 1 hr. at 4°C. Nuclei were washed twice in 200uL Dig150 Buffer to remove excess pA/G-MNase. Nuclei were resuspended in 100uL ice-cold Dig150 Buffer and cooled for at least 5 min. on ice.

CaCl_2_ was added to solution to 2mM to activate pA/G-MNase. Digestion was performed by incubating tubes on nutator for 2 hr. at 4°C. Digestion was quenched by addition of 100uL 2X Stop Buffer (340mM NaCl, 20mM EDTA, 4mM EDTA, 0.01% Digitonin, 100ug/mL RNAse A, 50ug/mL glycogen). Cleaved fragments were released into solution by incubating nuclei for 30 min. at 37°C. CUT&RUN DNA was isolated from solution via phenol:chloroform extraction, quantified via Qubit dsDNA assay (#Q32851, Invitrogen™, Eugene, OR, USA), and used for library preparation. Libraries were prepared using the NEBNext Ultra II FS DNA library preparation kit (#E7645 New England Biolabs, Ipswich, MA, USA). Briefly, 1-5ng of DNA was diluted to a final volume of 25uL in 0.1X TE. DNA was end-repaired with 5uL of End Repair Master Mix for 20 minutes at 20°C. Enzyme mix was inactivated for 1 hr. at 50°C and held at 4°C before proceeding to subsequent steps. End-repaired DNA was mixed with 1.25 uL of 1.5 uM Illumina Adapter and 15.5 uL of Ligation Master mix and incubated at 20°C for 15 min. 1uL of U-excision enzyme was added to each reaction and incubated at 37°C for 15 min. Adapter-ligated DNA fragments were isolated in 1X volume of AMPure XP beads (Beckman Coulter #A6381). DNA was resuspended in 12 uL 0.1X TE and 10.5 uL was used for indexing PCR. Input DNA was mixed with 2uL unique dual index oligos (#E6445 New England Biolabs, Ipswich, MA, USA) and 12.5uL NEBNext Q5 Ultra II PCR master mix. Indexing PCR was performed with the following thermocycler profile:

**Table 2.**
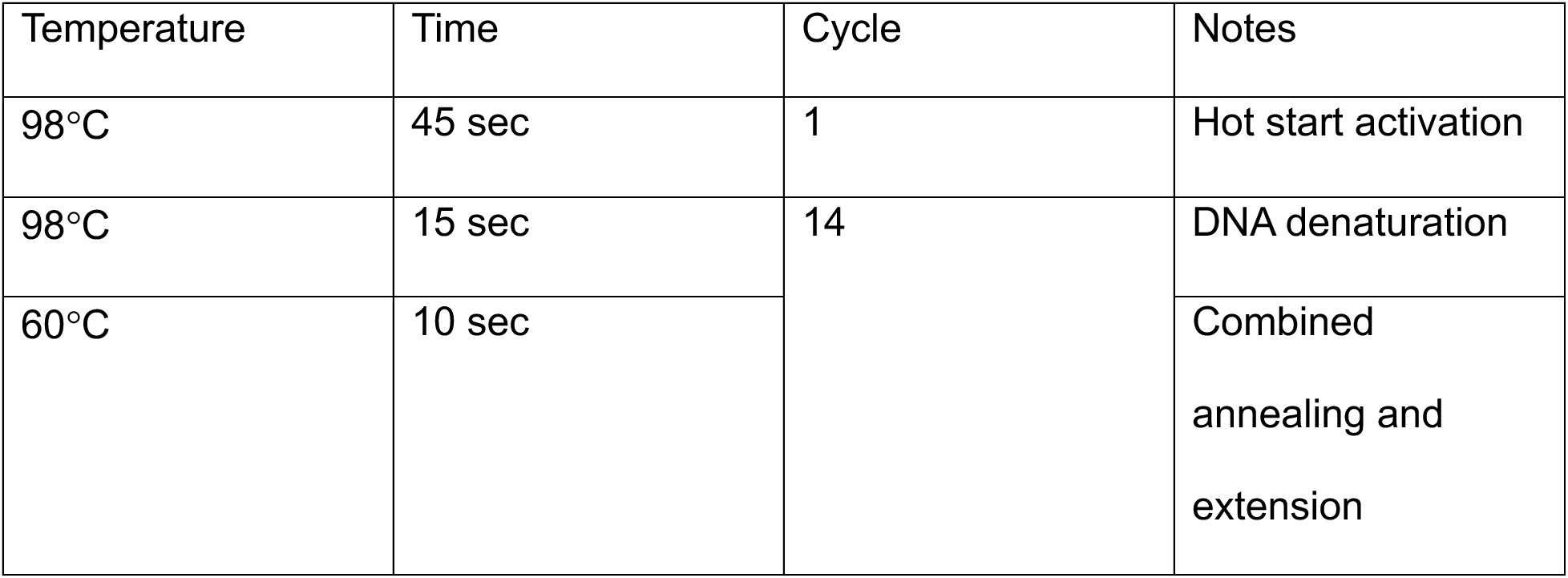
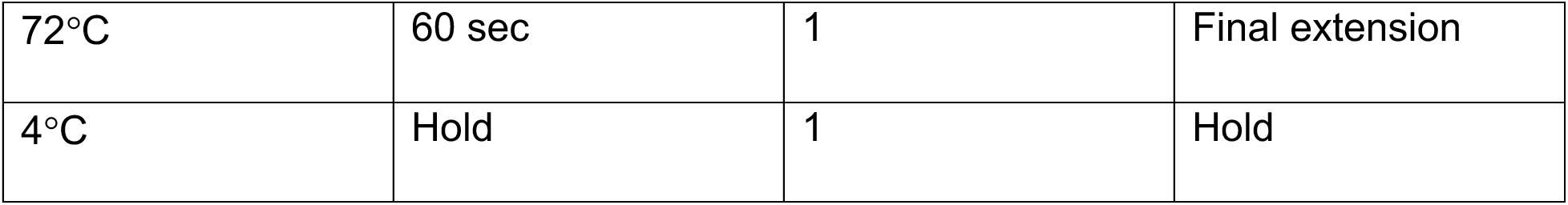
Thermocycler profile for CUT&RUN library preparation.

Libraries were sequenced on a NovaSeqX instrument to at least 10M paired-end reads per sample at the University of Wisconsin-Madison Biotechnology Center DNA Sequencing Core Facility (RRID:SCR_017759).

#### CUT&RUN-seq analysis

CUT&RUN paired-end reads were aligned to the mouse genome (mm39) using Bowtie2 version 2.5.4 with the following settings --local --very-sensitive --no-mixed --no- discordant --dovetail -I 10 -X 700 --no-unal. Aligned reads were filtered to exclude reads overlapping mm39 blacklist regions or with q score lower than 20 using samtools version 1.2. Coverage files were generated for each replicate using deeptools version 3.5.6 bamcoverage command with the following settings --binsize 20 -- effectiveGenomeSize 2654621783 --normalizeUsing RPKM -of bigwig. Average bigwig coverage files were generated for each group using deeptools bigwigaverage command with the following settings --binsize 20. H3K27me3 peaks were called for each replicate using the H3K27me3 aligned reads against the matched IgG control reads in MACS3 [67]. Consensus peaks were defined for each group using the MSPC workflow [68,69] with the following settings -r bio -s 1e-8 -w 1e-4 -g 1e-6 -p parser_config.json -- excludeHeader true. Read alignment and filtering, coverage file generation, and peak calling were done using a custom workflow on the University of Wisconsin Center for High Throughput Computing servers [70]. H3K27me3 signal was quantified over all H3K27me3 consensus peaks and subjected to differential enrichment analysis using DiffBind version 3.16.0 with significance threshold at FDR < 0.05. Differential peaks were analyzed for co-localization with naive H3K27me3 peaks using the findOverlaps command IRanges package (v2.44.0) [71]. H3K27me3 peaks were annotated using ChIPseeker v3.22 [72,73]. Briefly, peaks were assigned to genes if they co-localized with either the promoter, which we defined as -3000bp to +200bp of the transcription start site (TSS), or within the gene body. For analysis of H3K27me3 across TSS (Fig. 4C), counting windows were defined as -200 to +200bp of the TSS of the 241 network excitability genes. Number of H3K27me3 CUT&RUN fragments overlapping the 400bp TSS counting window were calculated for each sample using the featureCounts command of the Rsubread package (v2.20.0) [74]. Raw counts were then normalized to FPKM (fragments per kilobase million) based on the following formula:

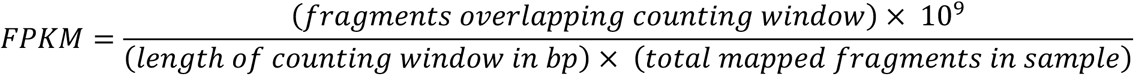

For CUT&RUN heatmaps and metaplots, plotting matrices were generated using deepTools version 3.5.6 computeMatrix command. Heatmap and metaplot matrices, and genome browser style plots were plotted in R (v.4.4.3) with custom code [75] utilizing the plotgardener package [76].

### Transcriptomics

#### RNA isolation and sequencing

Total RNA was isolated from flash-frozen whole hippocampi using TRIzol™ reagent (#15596026, Invitrogen) according to the manufacturer’s protocol. RNA was DNase treated (#AM2238, Invitrogen) followed by clean-up using RNeasy Mini Kit (#74104, Qiagen, Hilden, Germany). RNA library preparation and paired-end sequencing were performed by Novogene (Davis, CA, USA).

#### RNA-seq analysis

RNA counts were calculated from RNA paired-end reads using salmon (v1.10.3) [77] against the GRCm39 mouse transcriptome (GENCODE M36 release) [78]. Gene expression was quantified and subjected to differential analysis using DESeq2 (v1.46.0) [79]. DESeq2 output was filtered to retain only genes with 5 TPM in at least 1 sample and adjusted p-values were recalculated. Genes which were excluded were categorized as undetected by RNA-seq. Differentially expressed genes were defined as having at least 1.2-fold change and p-adjusted < 0.05.

#### Ontology analysis

Ontology analyses of H3K27me3 target genes were performed against the KEGG [20], Reactome [21], Biocarta [22,23], and Hallmark [24] databases using Ontomancer [13] using the list of all genes annotated to at least 1 H3K27me3 peak in our dataset as background. Ontology results were summarized using the ontomancer network analysis script according to Hoffman et al. [13] with resolution = 1 and beta = 6.

#### snRNA-seq meta-analysis

For meta-analysis of single-nuclei RNA-seq data from Hoffman et al. [13], network excitability gene score was calculated per nucleus using the scanpy (v1.12) score_genes function. For pseodobulk and differential expression analysis of neuronal cells, expression was pseudobulked across the following cell subcategories – DG (dentate granule cells), Pyramidal cells (CA1, CA2, CA3), Other excitatory neurons (Cajal-Retzius cells, Mossy cells), and Inhibhitory cells. Pseudobulked count matrices were used for differential expression analysis using DESeq2 (v1.46.0) [79]. DESeq2 output was filtered to retain only genes with 5 TPM in all samples of at least 1 condition and adjusted p-values were recalculated. Differentially expressed genes were defined as having p-adjusted < 0.1.

### Western Blot

Total protein lysate was prepared from flash-frozen hippocampi by lysing in Radioimmunoprecipitation Assay Buffer (RIPA: 50mM Tris, 150mM NaCl, 1% nonidet P- 40, 0.5% sodium deoxycholate, 0.1% SDS) with mammalian protease and phosphatase inhibitor (1:100; #PPC2020 Sigma-Aldrich). Tissue was homogenized in lysis buffer by probe sonication (Fisher Scientific, Sonic Dismembrator, Model 100, Hampton, NH) on power 3 for two rounds, with 10 pulses per round. Cell debris was pelleted by centrifugation at 16000xg at 4°C and the supernatant containing total protein lysate was collected. Total protein concentration was quantified using the DC Protein Assay (Bio-Rad, Hercules, CA) and stored at -80°C. For Western Blot, 5x SDS loading buffer (0.5mM Tris, 10% SDS, 50% glycerol, 10mM EDTA, 1% beta-mercaptoethanol) was added to each sample to reach a 1x final concentration. Extracts in loading buffer were boiled at 95°C on a heat block for 5 minutes and stored for up to 1 month at -20°C until run on an acrylamide gel. Protein extracts in loading buffer were loaded at 25ug per lane and resolved by electrophoresis in hand-poured acrylamide gels with a 5% acrylamide stacking layer (125mM Tris pH 6.8, 0.01% ammonium persulfate, 0.01% SDS, 0.01% TEMED) and 12% acrylamide separating layer (375mM Tris pH8.8, 12% acrylamide, 0.015% ammonium persulfate, 0.015% SDS, 0.08% TEMED). Gels were transferred to polyvinyl difluoride membranes (PVDF; Millipore, Bedford, MA) in Tris-glycine transfer buffer (20mM Tris, 1.5M glycine, 20% methanol). Membranes were blocked with 5% bovine serum albumin (Fisher Scientific, Fair Lawn, NJ) diluted in low-salt Tris-buffered saline (w/w TBST; 20mM Tris pH7.5, 150mM NaCl, 0.1% Tween-20) with mammalian protease and phosphatase inhibitor (1:1000; #PPC2020 Sigma-Aldrich) for 1 hour at room temperature. Primary antibodies were diluted in the same blocking buffer and incubated with membranes overnight at 4°C. Antibodies include: H2AK119ub (1:1000; #8240 Cell Signaling Technology, Inc., Danvers, MA, USA), Actin (1;10000; #MAB-1501 Millipore). The following day, membranes were washed three times in 1X TBST and incubated with horseradish peroxidase-conjugated goat-anti-mouse or -rabbit secondary antibodies for 1 hour at room temperature (1:10000; #31430 or #31460 Invitrogen, Rockford, IL). Membranes were subsequently washed three times in TBST, and membranes were developed in SuperSignal West Femto ECL or ECL reagent (ThermoFisher, Waltham, MA). Bands were imaged using a ChemiDoc-It Imaging System (UVP Vision-Works, Upland, CA) and quantified in ImageJ. Band intensities were analyzed and plotted in R v4.4.3.

### Statistical analysis

For all statistical analysis – unless otherwise specified – p<0.05 corrected for multiple comparisons was considered statistically significant. All statistical analyses were performed in R v4.4.3. Distributions of RNA expression response were analyzed by Pearson’s Chi-squared test. H3K27me3 TSS occupancy and mean RNA z-score expression data were analyzed by Kruskal-Wallis rank sum test followed by pairwise comparisons using Wilcoxon rank sum test with continuity corrections with multiple comparisons correction by Benjamini-Hochberg. Seizure threshold data was analyzed by 2-way ANOVA with repeated measures followed by Tukey’s multiple comparisons test. Seizure frequency data was analyzed by Kruskal-Wallis rank sum test. Change in seizure threshold and seizure frequency correlations were analyzed by Pearson correlation. Western Blot data for H2AK199ub and Actin were analyzed by Welch’s t- test.

## Supporting information

S1 Data. H3K27me3 DiffBind counts and results table

S2 Data. Enriched H3K27me3 peaks co-localization with naive H3K27me3 peaks analysis table

S3 Data. Normalized RNA counts and raw DESeq2 results

S4 Data. Re-calculated DESeq2 results after gene filtering

S5 Data. Enriched H3K27me3 peak annotation to gene table

S6 Data. Genes with no change and methylated ontomancer output

S7 Data. Genes induced and methylated ontomancer output

S8 Data. Genes induced and unmethylated ontomancer output

S9 Data. Genes repressed and methylated ontomancer output

S10 Data. Genes repressed and unmethylated ontomancer output

S11 Data. snRNA-seq meta-analysis network excitability gene score

S12 Data. snRNA-seq pseudobulk DESeq2 files

S13 Data. H3K27me3 signal quantification at TSS

S14 Data. RNA expression matrix UNC1999

S15 Data. Seizure threshold and spontaneous seizure recording data

S16 Data. H2AK119 WB raw images and quantification

## Data and Materials Availability

Mouse bulk CUT&RUN and RNA-seq data will be made available on the Gene Expression Omnibus (GEO) database at the National Center for Biotechnology Information. Analysis scripts in R and python will be made available on github/dryad. All other data associated with this study are present in the paper or available in the Supplementary Data files or upon request by contacting the corresponding author.

## Supplementary Materials

**S1 Fig.**
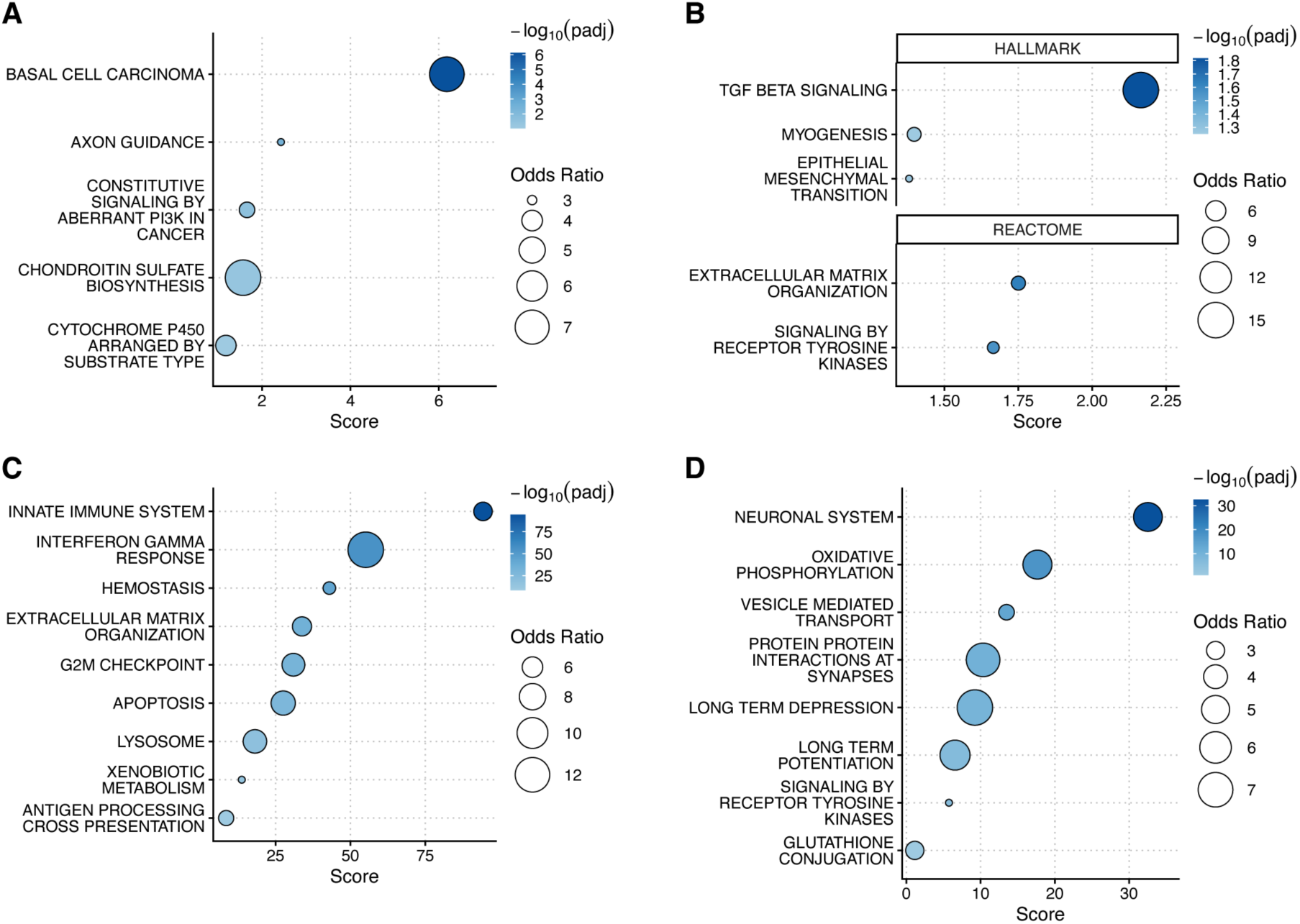
Summarized ontology analyses for different gene expression clusters. **(A)** H3K27me3 enriched genes with no change in RNA expression. **(B)** Induced genes with H3K27me3 enrichment. **(C)** Induced genes with no change in H3K27me3. **(D)** Repressed genes with no change in H3K27me3. Color is scaled to -log_10_(padj) value. Size is called to Odds Ratio.

**S2 Fig.**
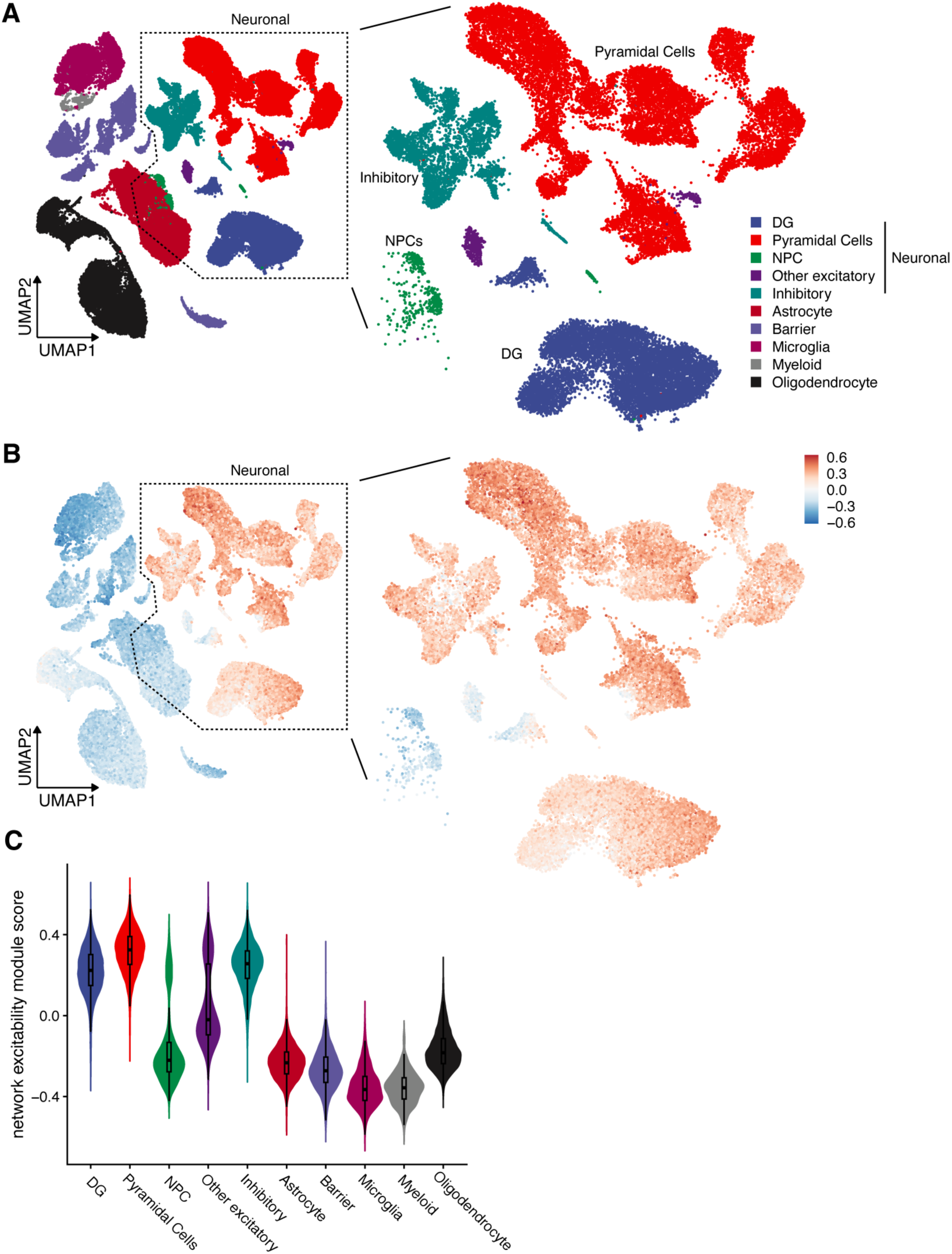
H3K27me3 target genes controlling network excitability are enriched in neuronal cells. **(A)** UMAP of re-annotated nuclei from a previously generated hippocampal snRNA-seq dataset in a mouse pilocarpine model of SE [13]. **(B-C)** Relative expression of the set of 241 H3K27me3 target network excitability gene was calculated across all nuclei. H3K27me3 target network excitability genes are enriched in DG, Pyramidal Cells, Other excitatory and Inhibitory neurons compared to non-neuronal cells and neural progenitor cells (NPCs). Violin and boxplots show median gene score and interquartile range per cell type.

**S3 Fig.**
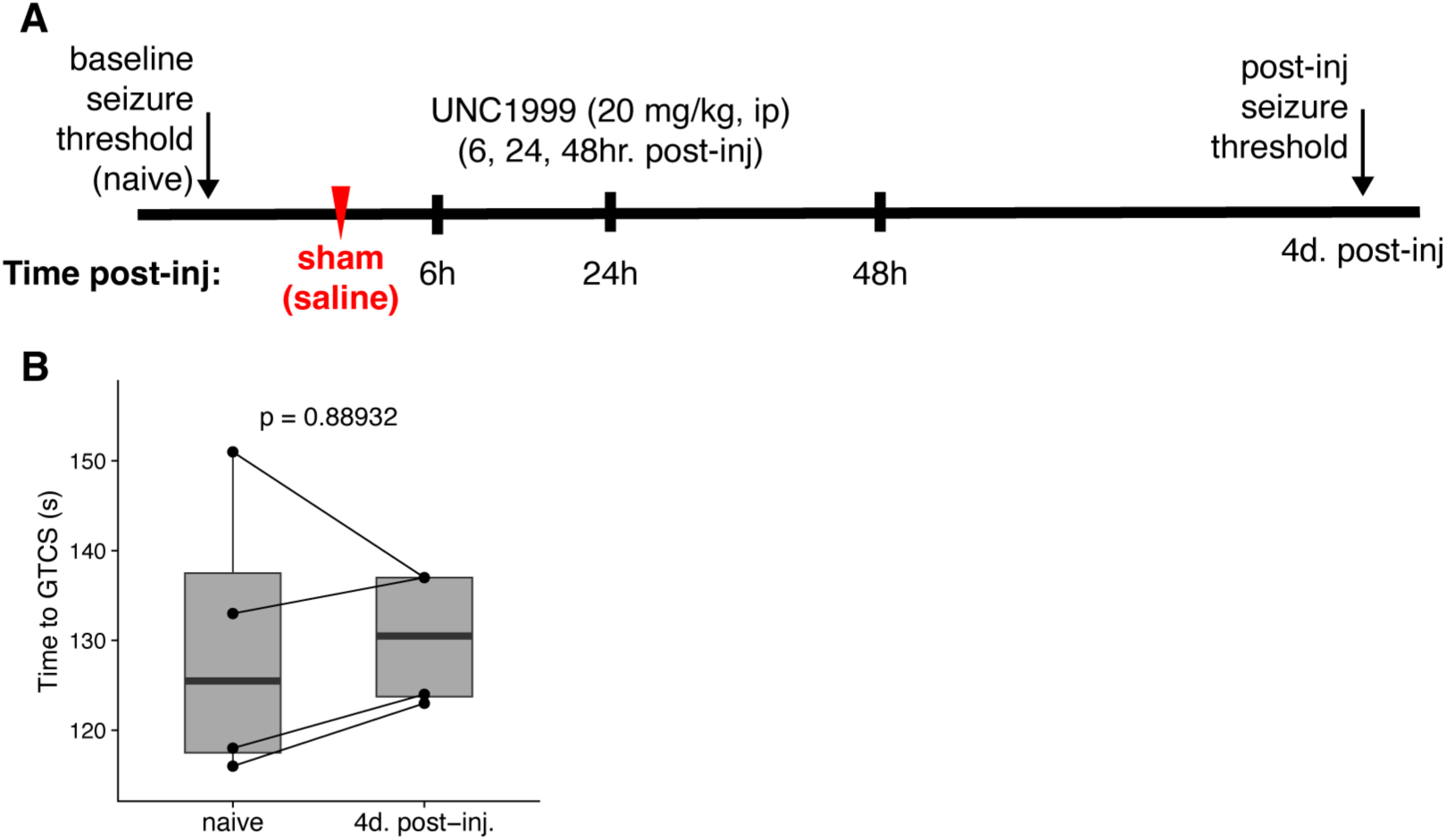
UNC1999 treatment does not increase network excitability in the absence of SE. **(A)** Naive animals were treated with UNC1999 after sham SE induction (see methods). For each mouse, seizure threshold was recorded prior to mock SE induction to determine naive baseline threshold and again 2 days after the final UNC1999 dose. **(B)** There was no difference in seizure threshold between baseline and post-treatment (Wilcoxon rank sum test). Lines connecting points indicate repeated measures for an individual mouse.

**S4 Fig.**
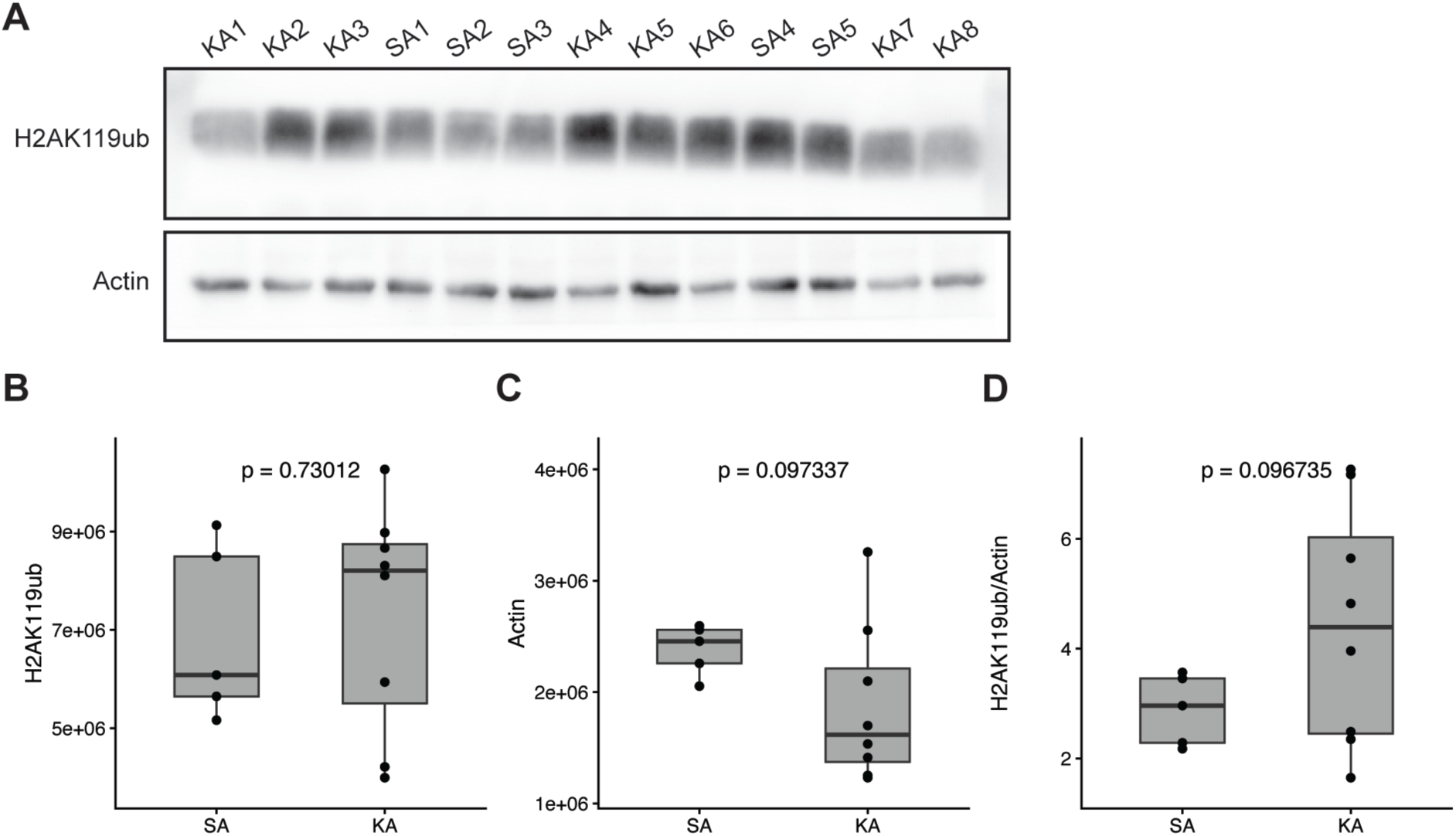
H2AK119ub is not significantly changed in the hippocampus post-SE. **(A)** Western blots showing H2AK119ub levels in hippocampal tissue of naive (n=5) and 4d. post-SE (n=8) mice. Actin was used as a loading control. Relative quantification by densitometry showed no significant difference (Welch’s t-test) in **(B)** raw H2AK119ub and **(C)** raw Actin signal between naive or post-SE samples. **(D)** H2AK119ub normalized to Actin levels show no significant difference (Welch’s t-test) between naive and post-SE samples. Boxplots show the median and interquartile range per group.

**S5 Fig.**
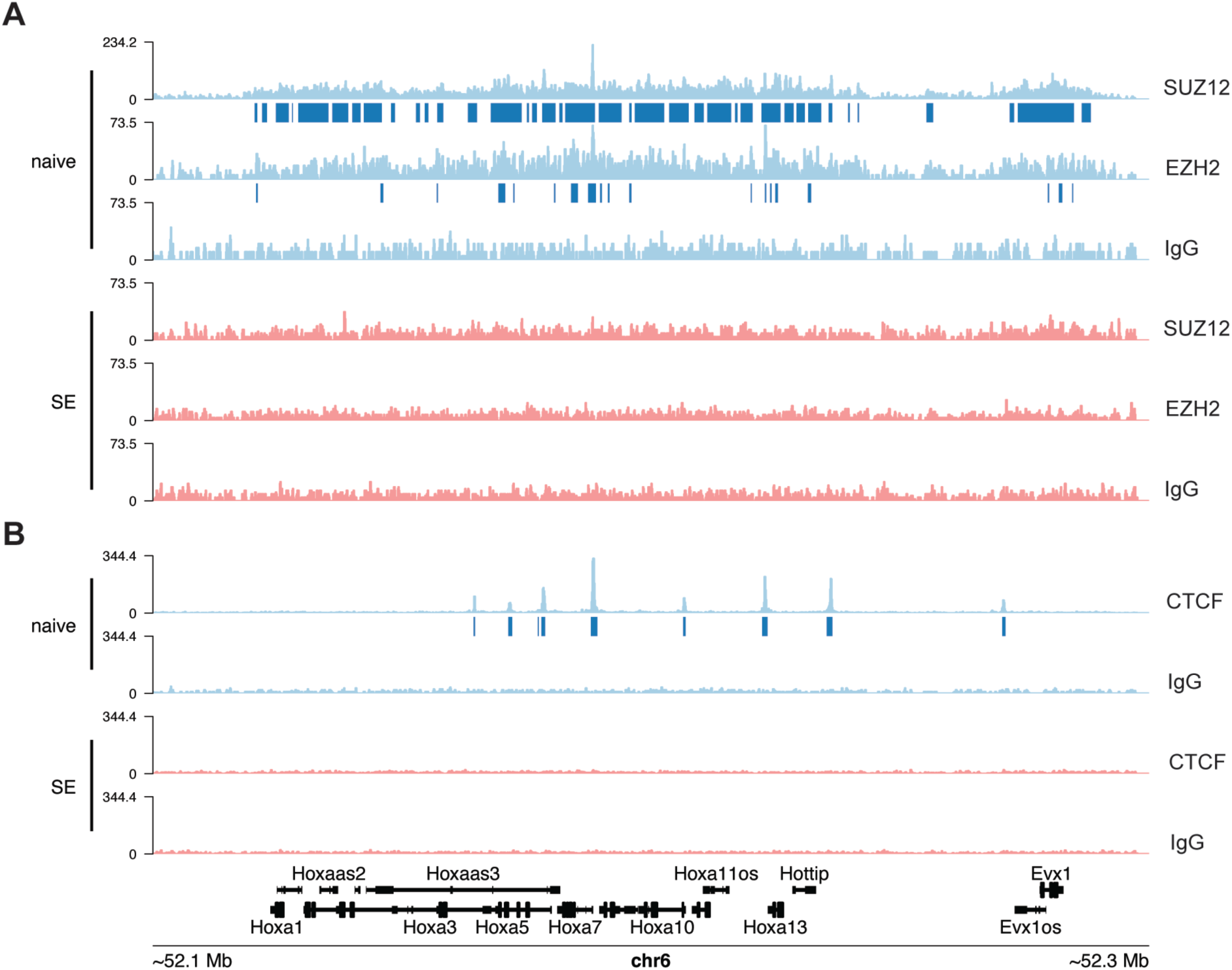
Post-SE hippocampi are non-permissive to transcription factor/co-factor CUT&RUN. **(A)** SUZ12, EZH2, and **(B)** CTCF CUT&RUN coverage (FPKM) plotted across the Hox gene cluster. Peaks are plotted as rectangles underneath the coverage plots. CUT&RUN successfully identifies SUZ12, EZH2, and CTCF peaks in naive hippocampi but not in post-SE samples. Control IgG coverage is also plotted for comparison. Scale annotations are plotted on the y-axis.

**S1 Data.** H3K27me3 DiffBind counts and results table

**S2 Data.** Enriched H3K27me3 peaks co-localization with naive H3K27me3 peaks analysis table

**S3 Data.** Normalized RNA counts and raw DESeq2 results

**S4 Data.** Re-calculated DESeq2 results after gene filtering

**S5 Data.** Enriched H3K27me3 peak annotation to gene table

**S6 Data.** Genes with no change and methylated ontomancer output

**S7 Data.** Genes induced and methylated ontomancer output

**S8 Data**. Genes induced and unmethylated ontomancer output

**S9 Data.** Genes repressed and methylated ontomancer output

**S10 Data**. Genes repressed and unmethylated ontomancer output

**S11 Data.** snRNA-seq meta-analysis network excitability gene score

**S12 Data.** snRNA-seq pseudobulk DESeq2 files

**S13 Data.** H3K27me3 signal quantification at TSS

**S14 Data.** RNA expression matrix UNC1999

**S15 Data**. Seizure threshold and spontaneous seizure recording data

**S16 Data.** H2AK119 WB raw images and quantification

