## Supplementary material for "H3K27 trimethylation maintains baseline network excitability in epilepsy": S16 Data. H2AK119 WB raw images and quantification: H2AK119ub_WB_analysis.Rmd

````
---
title: "H2AK119_WB_analysis"
author: "JEspina"
date: "2026-07-17"
output: html_document
---

```{r setup, include=FALSE}
knitr::opts_chunk$set(echo = TRUE)
```

RMD for plotting WB quantification of H2AK119ub

```{r}
library(ggplot2)
library(dplyr)
library(forcats)
library(readr)
library(stringr)
```

```{r}
## Read in sample order
sample_order <- read_tsv(file="../sample_order.txt") %>% pull(sample)

## Read in results
H2AK119 <- read.csv(file = "H2AK119ub/H2AK119ub_ROI_quant.csv") %>% 
  select(-X) %>% 
  mutate(sample = sample_order) %>%
  select(sample,signal) %>%
  mutate(condition = str_sub(sample, start=1, end=2))

H2AK119$condition <- as.factor(H2AK119$condition) %>% fct_relevel("SA")

Actin <- read.csv(file="Actin_for_H2AK119ub/Actin_quantification.csv") %>% 
  select(-X) %>% 
  mutate(sample = sample_order) %>%
  select(sample,signal) %>%
  mutate(condition = str_sub(sample, start=1, end=2))

Actin$condition <- as.factor(Actin$condition) %>% fct_relevel("SA")

```
```{r}
## Calculate raw
H2AK119_test <- t.test(
  H2AK119$signal[H2AK119$condition == "SA"],
  H2AK119$signal[H2AK119$condition == "KA"]
)

Actin_test <- t.test(
  Actin$signal[Actin$condition == "SA"],
  Actin$signal[Actin$condition == "KA"]
)
```

```{r}
## Plot raw
H2AK119_plot <- ggplot(data = H2AK119,
       aes(x = condition, y = signal)) + 
  geom_boxplot(width = 0.5, fill = "darkgray") + 
  geom_point(size = 1) +
  ## annotate plot
  geom_text(data = tibble(x = 1.5,
                          y = c(10E06,10E06),
                          label_text = paste0("p = ", formatC(H2AK119_test$p.value,digits = 5))
                          ),
            aes(x=x, y=y, label = label_text),
            inherit.aes = FALSE) +
  
  ## adjust plot appearance
  scale_y_continuous(expand = expansion(mult = c(0.1,0.1))) +
  theme_classic(base_size = 7) +
  theme(aspect.ratio=1,
        legend.key.size = unit(3,"mm"), 
        legend.position = "right",
        legend.justification = "top"
        ) +
  coord_cartesian(clip = "off") + 
  ## adjust plot labels
  labs(
    x = "",
    y = "H2AK119ub"
  )

H2AK119_plot

Actin_plot <- ggplot(data = Actin,
       aes(x = condition, y = signal)) + 
  geom_boxplot(width = 0.5, fill = "darkgray") + 
  geom_point(size = 1) +
  ## annotate plot
  geom_text(data = tibble(x = 1.5,
                          y = c(4E06,4E06),
                          label_text = paste0("p = ", formatC(Actin_test$p.value,digits = 5))
                          ),
            aes(x=x, y=y, label = label_text),
            inherit.aes = FALSE) +
  
  ## adjust plot appearance
  scale_y_continuous(expand = expansion(mult = c(0.1,0.1))) +
  theme_classic(base_size = 7) +
  theme(aspect.ratio=1,
        legend.key.size = unit(3,"mm"), 
        legend.position = "right",
        legend.justification = "top"
        ) +
  coord_cartesian(clip = "off") + 
  ## adjust plot labels
  labs(
    x = "",
    y = "Actin"
  )

Actin_plot

```


```{r}
## Calculate normalized
normalized <- data.frame(
  sample = Actin$sample,
  norm = H2AK119$signal/Actin$signal,
  condition = Actin$condition
)

normalized$condition <- normalized$condition %>% as.factor() %>% fct_relevel("SA")

t_test <- t.test(
  normalized$norm[normalized$condition == "SA"],
  normalized$norm[normalized$condition == "KA"]
  )

wilcox.test(
  normalized$norm[normalized$condition == "SA"],
  normalized$norm[normalized$condition == "KA"]
  )

```
```{r}
norm_plot <- ggplot(data = normalized,
       aes(x = condition, y = norm)) + 
  geom_boxplot(width = 0.5, fill = "darkgray") + 
  geom_point(size = 1) +
  ## annotate plot
  geom_text(data = tibble(x = 1.5,
                          y = c(7, 7),
                          label_text = paste0("p = ", formatC(t_test$p.value,digits = 5))
                          ),
            aes(x=x, y=y, label = label_text),
            inherit.aes = FALSE) +
  
  ## adjust plot appearance
  scale_y_continuous(expand = expansion(mult = c(0.1,0.1))) +
  theme_classic(base_size = 7) +
  theme(aspect.ratio=1,
        legend.key.size = unit(3,"mm"), 
        legend.position = "right",
        legend.justification = "top"
        ) +
  coord_cartesian(clip = "off") + 
  ## adjust plot labels
  labs(
    x = "",
    y = "H2AK119ub/Actin"
  )

norm_plot
```

```{r}
## plot on plotgardener
library(ggpubr)

a <- H2AK119_plot
b <- Actin_plot
c <- norm_plot

plots <- ggarrange(a,b,c, nrow=1, ncol=3, align="v")
plots
```
```{r}
library(plotgardener)
units = pgParams(default.units="cm")

## Plot figures on page
## Set page dimensions
page_width=20
page_height=6

pdf(file = "H2AK119ub_WB_plots.pdf", width = page_width/2.54, height = page_height/2.54, useDingbats = FALSE)

## Plot on plotgardener
# generate plotGardener page
pageCreate(width = page_width, height = page_height, default.units = "cm", showGuides = FALSE)

# set reference axes
ref_x = 0.5
ref_y = 0.5

plotGG(
  plot = plots,
  x = ref_x, y = ref_y,
  width = 16, height = 5,
  just = c("left", "top"),
  params = units
)

dev.off()
```
````
