## Supplementary figures and images for "H3K27 trimethylation maintains baseline network excitability in epilepsy"

### 20220622 Actin_H2AK119ub-blot_femto_5s.TIF

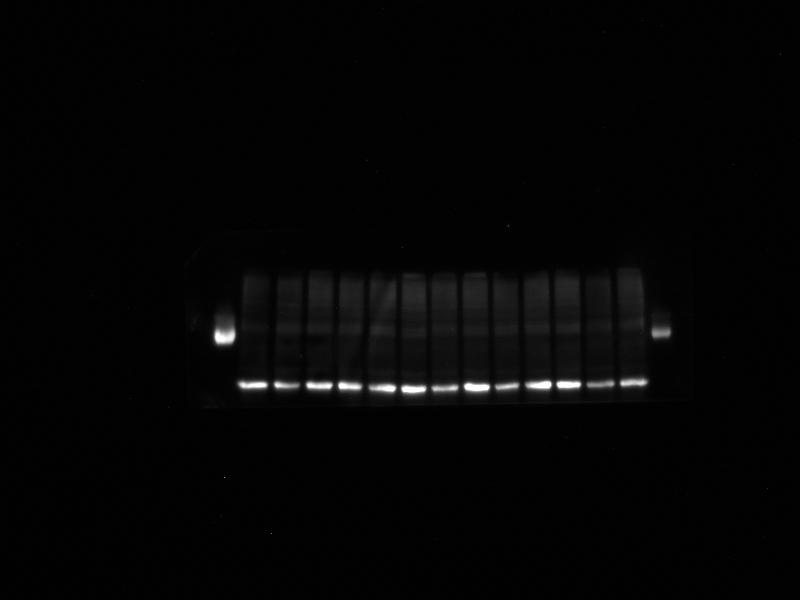

### 20220622 Actin_H2AK119ub-blot_femto_5s_inverted.tif

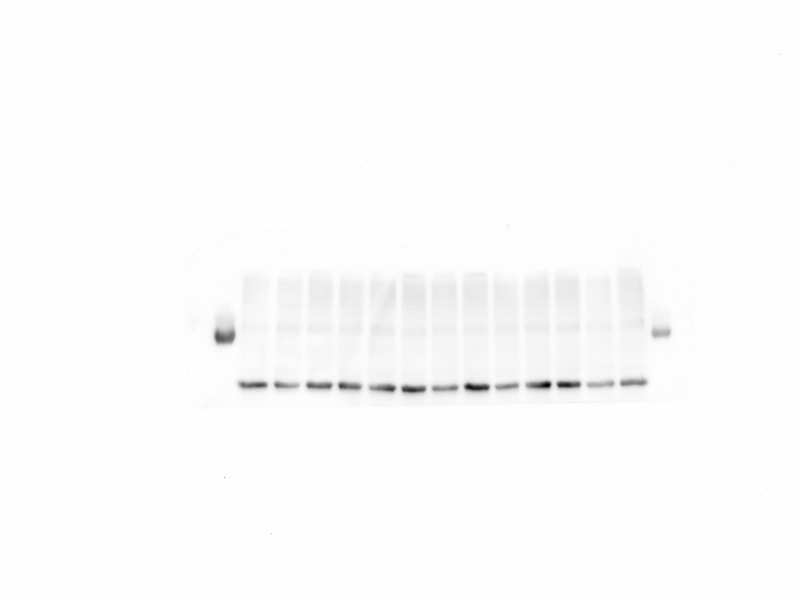

### 20220622 Actin_H2AK119ub-blot_femto_5s_wl.TIF

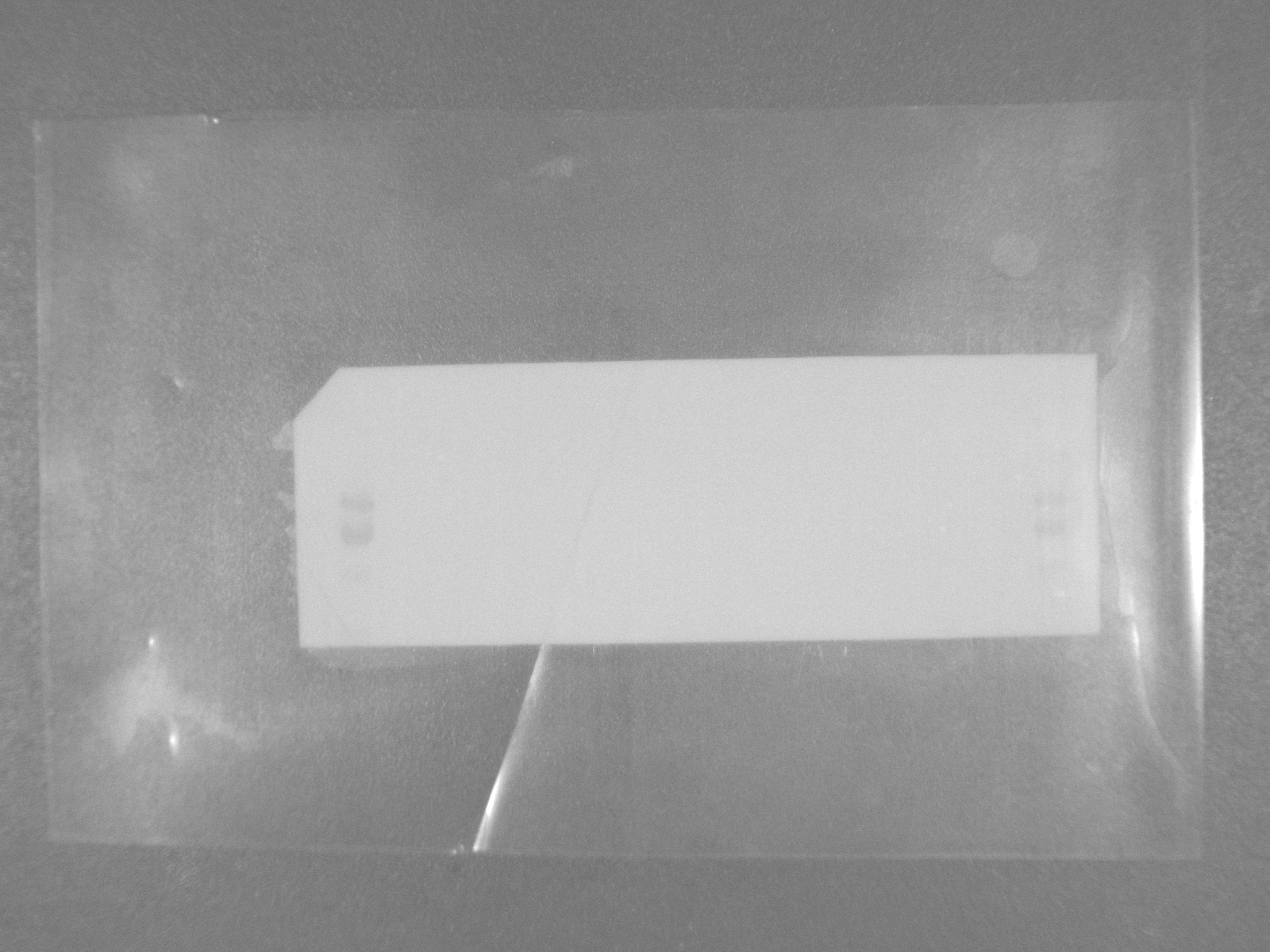

### 20220622 H2AK119ub_femto_50ms.TIF

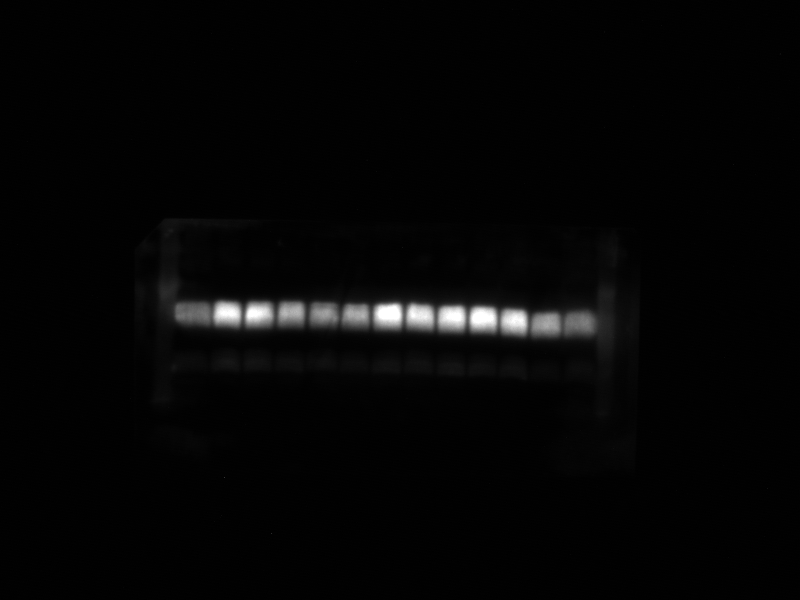

### 20220622 H2AK119ub_femto_50ms_inverted.TIF

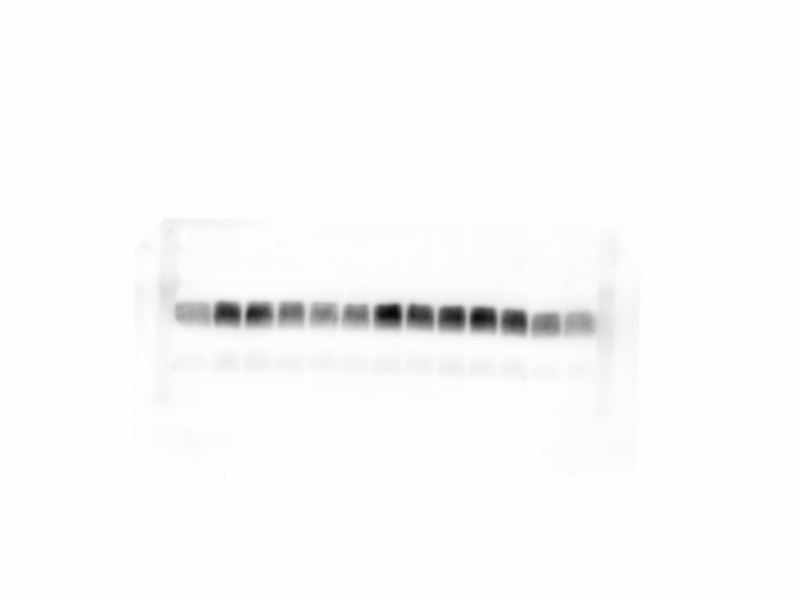

### 20220622 H2AK119ub_femto_50ms_wl.TIF

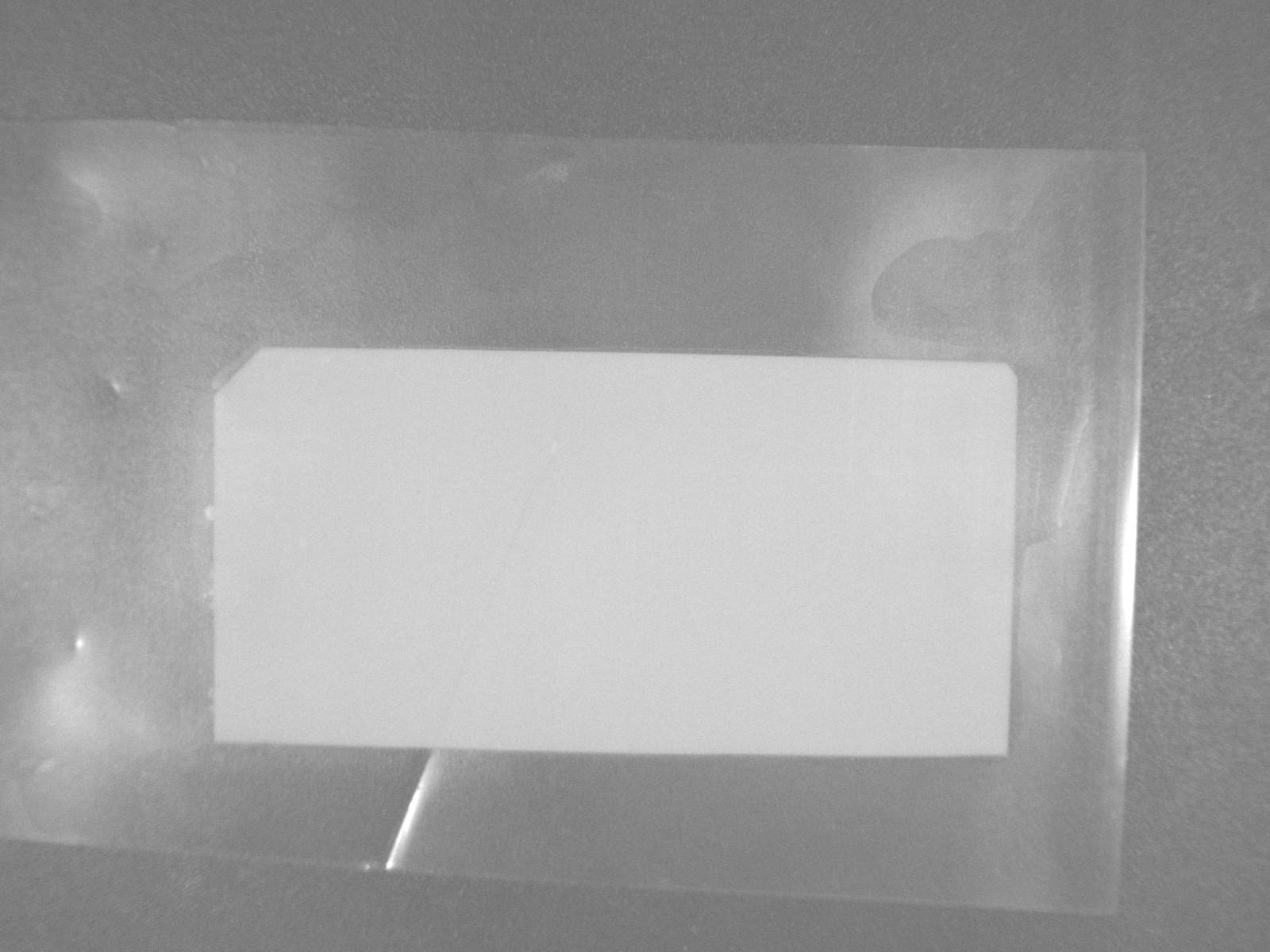
